# *Ace* and *ace*-like genes of invasive redlegged earth mite: Copy number variation, target-site mutations, and their associations with organophosphate insensitivity

**DOI:** 10.1101/2022.11.02.514930

**Authors:** Joshua A Thia, Paul A Umina, Ary A Hoffmann

## Abstract

**BACKGROUND:** Invasive Australian populations of redlegged earth mite, *Halotydeus destructor* (Tucker), are evolving increasing organophosphate resistance. In addition to the canonical *ace* gene, the target gene of organophosphates, the *H. destructor* genome contains many radiated *ace*-like genes that vary in copy number and amino acid sequence. In this work, we characterise copy number and target-site mutation variation at the canonical *ace* and *ace*-like genes and test for potential associations with organophosphate insensitivity. This was achieved through comparisons of whole-genome pool-seq data from alive and dead mites following organophosphate exposure.

**RESULTS:** A combination of increased copy number and target-site mutations at the canonical *ace* was associated with organophosphate insensitivity in *H. destructor*. Resistant populations were segregating for G119S, A201S, F331Y at the canonical *ace*. A subset of populations also had copy numbers of canonical *ace* >2, which potentially helps over-express proteins carrying these target-site mutations. Haplotypes possessing different copy numbers and target-site mutations of the canonical *ace* gene may be under selection across *H. destructor* populations. We also detected some evidence that increases in copy number of radiated *ace*-like genes are associated with organophosphate insensitivity, which might suggest potential roles in sequestration or breakdown of organophosphates.

**CONCLUSION:** Different combinations of target-site mutations and (or) copy number variation in the canonical *ace* and *ace*-like genes may provide non-convergent ways for *H. destructor* to respond to organophosphate selection. However, these changes may only play a partial role in organophosphate insensitivity, which appears to have a polygenic architecture.

## Introduction

Pesticide resistance in arthropod pests has been extensively studied to understand the genetic basis of adaptation to toxins. Many pesticides exert their toxicity through very specific structural interactions with a target protein (e.g., Russell *et al*., 2004; Davies *et al*., 2007; Crossthwaite *et al*., 2014). Such precise biomolecular interactions commonly produce convergent evolution of the same mutations in the target gene (‘target-site mutations’) that confer pesticide resistance (Foster et al., 1996; Khajehali et al., 2010; Martinez-Torres, Devonshire, & Williamson, 1997). This observed convergence in target-site mutations is striking because it occurs across diverse taxonomic levels. For example, the target gene of organophosphate pesticides is the enzyme acetylcholinesterase (*ace*), which is responsible for breaking down to acetylcholine in the synapse (Russell et al., 2004). Organophosphorus pesticides inhibit this action by competitively binding to *ace*, preventing acetylcholine breakdown, leading to uncontrolled firing at the synapse and ultimately death (Oakeshott et al., 2005). Many arthropod pests have evolved the same mutations that disrupt inhibition of *ace* by organophosphates, a convergence on the exact same genetic mechanism underpinning resistance (reviewed in: Fournier, 2005; Feyereisen*et al*., 2015).

However, the genetic basis of pesticide resistance can be more complex than mutational changes in target sites. Duplications of target genes can contribute to resistance by increasing gene dosage, elevating the number of target protein molecules that must be bound by a pesticide to achieve mortality (Kwon, Clarkt, & Lee, 2010; Deok Ho Kwon, Choi, Je, & Lee, 2012). Target gene duplications can also propagate haplotypes carrying target-site mutations in the genome, increasing the proportion of target protein molecules carrying pesticide resistance mutations (Carvalho et al., 2012; Kwon et al., 2010). Alternatively, increased expression of duplicated target genes may be beneficial because a comparable level of enzymatic activity is retained despite reduced functionality of resistant proteins (Deok Ho Kwon et al., 2012). Target-site mutations can, however, come with considerable fitness costs (Freeman, San Miguel, & Scott, 2021; Langmüller et al., 2020). Other genes unrelated to target-sites can also contribute to pesticide resistance, including genes involved in detoxification processes (Crossley, Chen, Groves, & Schoville, 2017; Devonshire & Moores, 1982; Walsh et al., 2018), and (or) genes that help reduce cuticle penetration by pesticides (Crossley et al., 2017; Hsu et al., 2016).

In this study, we focus on the redlegged earth mite, *Halotydeus destructor* (Tucker 1925; Penthaleidae, Trombidiformes), an agricultural pest of Australian grain and pasture crops. This mite invaded the state of Western Australia from South Africa around 1917 and spread throughout the southern regions of Australia during the 1920s (Newman, 1925; reviewed by Ridsdill-Smith, 1997). It is a winter-active pest, which diapauses as eggs during the Australian summer (October–March), emerges in autumn (April–May), and typically undergoes its active life stage through to late spring (April–November) (Ridsdill-Smith, 1997). Control of this mite for many decades has been largely achieved through the application of pyrethroid and organophosphate pesticides (Arthur et al., 2021; Paul A. Umina, McDonald, Maino, Edwards, & Hoffmann, 2019). Low-level tolerance to these pesticides was initially observed in the 1990s and early 2000s (Hoffmann, Porter, & Kovacs, 1997; Robinson & Hoffmann, 2000), with high-level pyrethroid resistance first detected in field populations in 2006 (Paul A Umina, 2007) and organophosphate resistance first detected in 2014 (Umina et al. 2017).

Whilst the molecular mechanism of pyrethroid resistance in *H. destructor* has mostly been linked to target-site mutations (Cheng, Umina, Lee, & Hoffmann, 2019; Edwards et al., 2018; Yang et al., 2020), the molecular mechanism of organophosphate resistance has been difficult to elucidate. We recently assembled a draft genome for *H. destructor*, which helped reveal a large radiation event in *ace*-like genes (Thia et al., 2023). Our previous comparative analyses showed that such a radiation of *ace*-like is highly unusual among other arthropod pests but has also been observed in *Tetranychus urticae*, the two-spotted mite, another problematic trombidiform mite pest (Thia et al., 2023). There are some 28 distinct genes, which we refer to as the canonical *ace* gene or *ace*-like genes in the *H. destructor* genome. The protein products of these genes vary in size from 231–1096 amino acids in length; typically, *ace* genes are around 600 amino acids long and the standard reference from *Torpedo californica* is 596 amino acids long (GenBank: X03439.1).

We had previously reported the presence of known organophosphate target-site mutations (G119S, A201S, and F331Y) in the canonical *ace* gene (Thia et al., 2023) that are segregating in resistant populations. However, the amino acid sequence variation in the radiated *ace*-like genes has yet to be formally characterised. We also suspect that the canonical *ace* and ace-like genes may vary in copy number (Thia et al., 2023). The exact functional roles of radiated *ace*-like genes in *H. destructor* are currently unknown. However, if some of these *ace*-like genes can bind organophosphates, they could help reduce inhibition of canonical *ace* activity by sequestering pesticide molecules (Kim, Cha, Jung, Kwon, & Lee, 2012).

Organophosphates may also select for mutations in esterases that allow them to hydrolyse organophosphates, reducing inhibition of canonical *ace* activity through metabolism of pesticide molecules (Newcomb et al., 1997). If any of these radiated *ace*-like genes can contribute to organophosphate insensitivity, there may be diverse ways in which they could combine with the canonical *ace* to influence the phenotypes of *H. destructor* (Figure 1).

**Figure 1.**
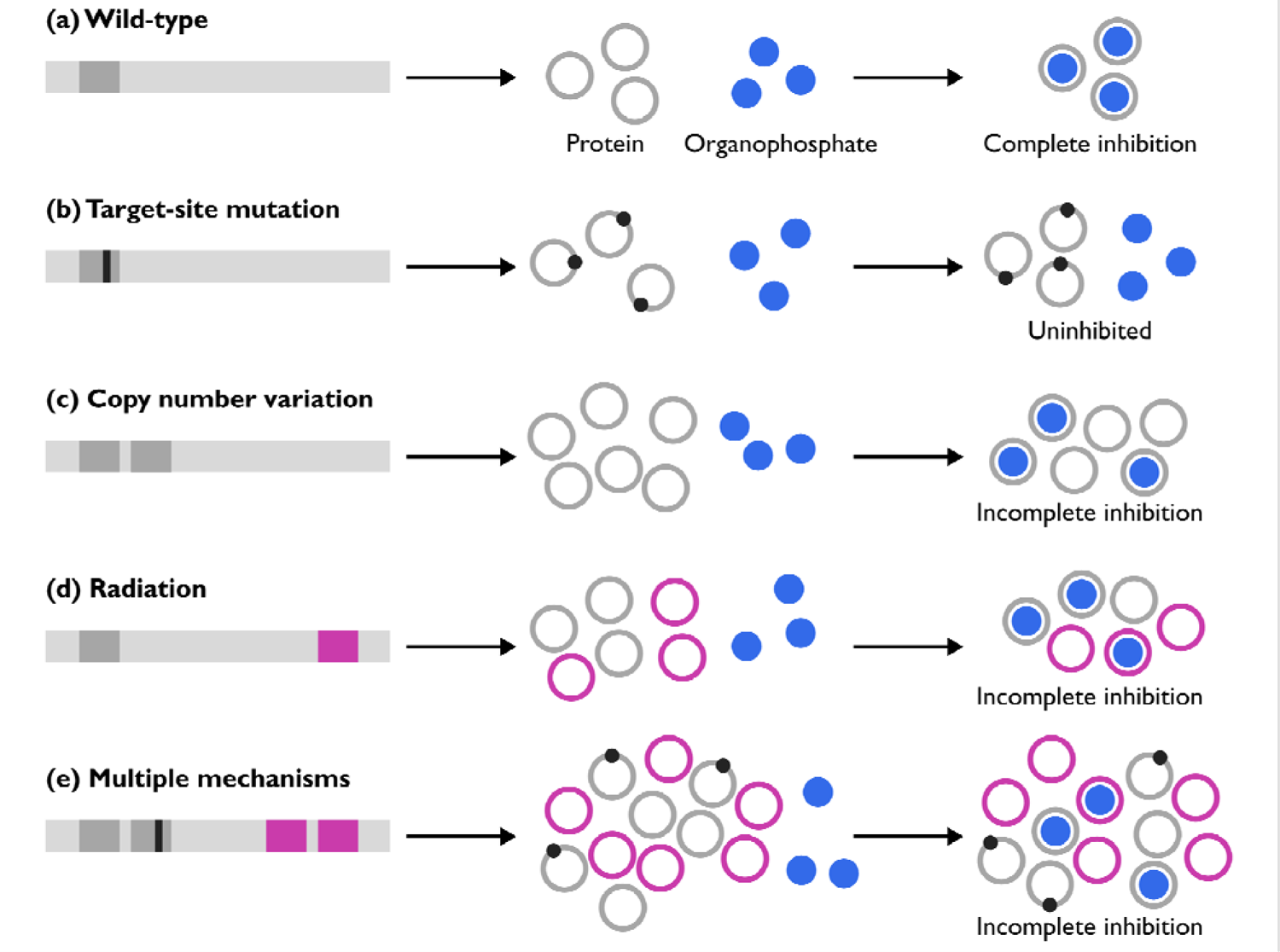
Schematic of hypothesized mechanisms for canonical *ace* and radiated *ace*-like genes to organophosphate insensitivity. All mechanisms reduce inhibition of the canonical *ace*. (a) The wild-type canonical *ace* gene produces protein that is sensitive to organophosphate inhibition. (b) Target-site mutations in the canonical *ace* gene allow production of protein th at is insensitive to organophosphate inhibition. (c) Increasing copy number of the canonical *ace* helps over-express the protein product, requiring more organophosphate molecules to achieve complete inhibition. (d) Radiated *ace* genes help sequester or breakdown organophosphate pesticides, reducing inhibition of canonical *ace* proteins. (e) Multiple mechanisms combine to reduce inhibition of the canonical *ace* proteins.

In this study, we characterised copy number variation and target-site amino acid variation in the *ace* and *ace*-like genes of *H. destructor* and test for associations with organophosphate insensitivity. We also explored the role of other non-*ace* genes in responses to organophosphates. We used a pool-seq approach to obtain whole-genome data from mites that were alive (insensitive) and dead (sensitive) following organophosphate exposure. These mites came from populations with different levels of evolved organophosphate resistance.

Our results are most consistent with multiple mechanism of resistance by which different combinations of target-site changes and (or) copy number variation in the canonical *ace* and *ace*-like genes may contribute to organophosphate insensitivity in *H. destructor*. However, these effects are variable across populations and they can be small, suggesting that organophosphate insensitivity often has a polygenic architecture that encompasses other (uncharacterized) mechanisms. The potential polygenicity of organophosphate insensitivity was evident from a lack of clear signals of selection in genomic outlier scans. These findings suggest non-convergent evolution of organophosphate resistance across populations.

## Methods

### Terminology and definitions

#### Organophosphate insensitivity

An individual (or population) is typically considered ‘resistant’ to a pesticide when it has evolved to survive exposure to high chemical doses. In contrast, ‘susceptible’ individuals (or populations) are those that have not evolved resistance and are killed at doses that otherwise ‘resistant’ individuals (or populations) would survive at. At a high exposure dose, the nomenclature ‘resistant’ versus ‘susceptible’ is intuitive but is more subjective when high doses cannot be tolerated, and (or) if a low exposure dose is used. We therefore use the term ‘organophosphate insensitivity’ in this study as a more general way of separating individuals exposed to pesticide from populations with different phenotypic means. Here, individuals within populations that could survive organophosphate exposure are considered ‘insensitive’, whereas those that were killed by exposure are considered ‘sensitive’. This definition allows us to refer to responses of phenotypically ‘resistant’ and ‘susceptible’ populations more generally.

#### *Ace* and *ace* -like genes

We use the term ‘ *ace*-like’ for annotated genes with similar sequences to *ace* that appear to have radiated in the *H. destructor* genome. These radiated *ace*-like genes are all divergent from the ‘canonical *ace*’ gene, the putative functional version of *ace* that is orthologous to those in other arthropods (Thia et al., 2023). Although we are yet to determine the functional characteristics of the radiated *ace*-like genes, we hypothesize some may contribute to organophosphate insensitivity through different potential mechanisms (Figure 1). We consider that the canonical *ace* gene and radiated *ace*-like genes exist as their own lineage. This is an important definition because we show in this study that each unique *ace* or *ace*-like gene can possess copy number variation within and among populations. In this work, we focus on the canonical *ace* gene and 18 focal *ace*-like genes that reside on large genomic contigs (see below). We refer to these genes by annotation codes, ‘HDE_X’, where ‘X’ is a unique gene identifier.

#### Biological material

*Halotydeus destructor* collected for this study came from industry projects surveying phenotypic resistance in the states of Western Australian and Victoria (Figure 2) (described in Thia et al., 2023). All Western Australia populations (Manjimup, Tambellup, Treeton, and Yongarillup) were found to be resistant to organophosphates. Victoria populations included two from Colbinabbin, one with evolved resistance to organophosphates (herein referred to as Colbinabbin Res) and the other susceptible to organophosphates (herein referred to as Colbinabbin Sus) (Thia, Cheng, Maino, Umina, & Hoffmann, 2022). A third Victorian population was collected from Wantirna South and was previously used to assemble the reference genome for *H. destructor* (Thia et al., 2023). Although not phenotypically screened in this study, mites from Wantirna South are known to be susceptible to organophosphates.

**Figure 2.**
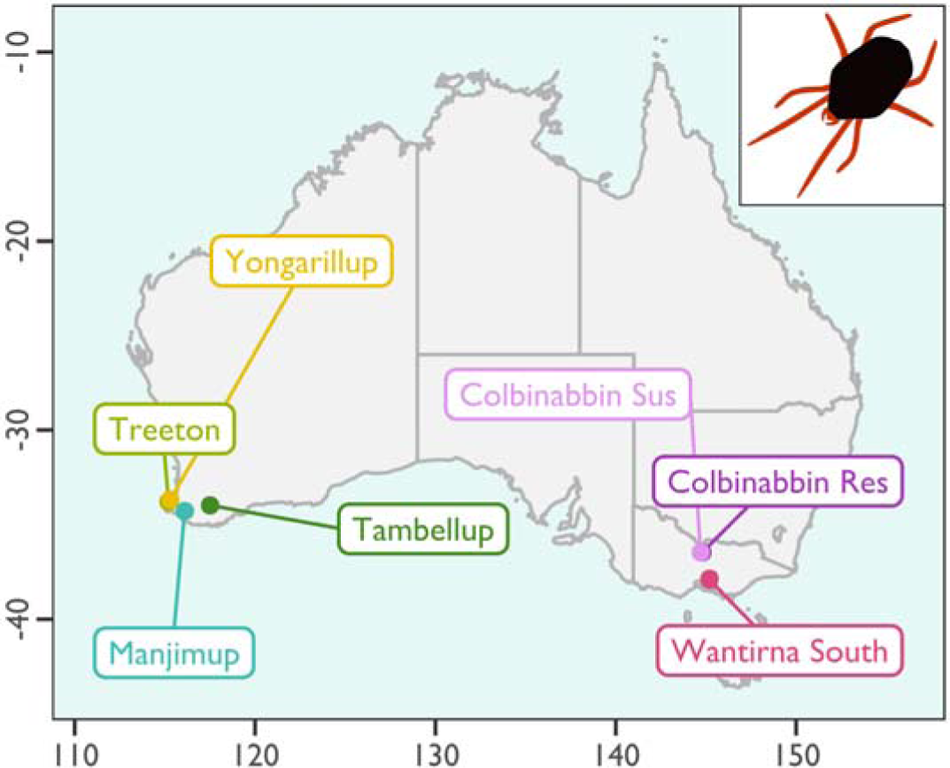
Map of Australia illustrating the locations of our studied populations

Collections of mites from phenotypic resistance surveys were exposed to either omethoate (Western Australia populations) or dimethoate (Victoria populations), two widely used organophosphates for mite control in Australia. The Western Australian mites used in this study were exposed to either 5.8 or 8.7 mgL^-1^ omethoate; 5.8 mgL^-1^ is a standard discriminating omethoate dose for resistance in *H. destructor* that kill phenotypically susceptible mites (Arthur et al., 2021). Colbinabbin Res mites were exposed to 34 mgL^-1^ of dimethoate, and Colbinabbin Sus mites were exposed to 3.4 mgL^-1^ dimethoate. These doses produce roughly 60% mortality (LD_60_ estimate) in their respective populations, and all mites from the Colbinabbin Sus population die at 34 mgL^-1^ (see Figure S1). Pesticide exposures were implemented using the coated vial method (Robinson & Hoffmann, 2000; Umina & Hoffmann, 1999). Briefly, mites were collected from the field and held at 4°C before exposure to pesticide. Pesticides were prepared by diluting stock solutions of 290 gL^-1^ omethoate (Le Mat® Bayer CropScience) or 400 gL^-1^ dimethoate (Dimethoate ADAMA) in distilled water with 0.01% polysorbate 80 (Tween^®^), and then diluted to make up the necessary doses. Pesticides were swirled in 30 mL plastic vials to coat and left to air-dry overnight. The following day, mites were placed into each vial along with a single vetch (*Vicia sativa*) leaf. Mites from Western Australia were incubated for 24 hours at 18°C, following Arthur et al. (2021). The Victorian mites were incubated for 24 hours at 11°C, following Thia, Cheng, Maino, Umina, & Hoffmann (2022), which produced mortality under thermal conditions more realistic to those in the field. Approximately 50 mites were tested at each dose. Mites were scored as alive (insensitive) and dead (sensitive), separated, placed into Eppendorf tubes with 100% ethanol, and stored at −20°C for molecular analysis. For some populations, we kept samples of untreated mites that were either unexposed to pesticides (Colbinabbin Res and Colbinabbin Sus) or were exposed to a dose that imposed 0% mortality (Manjimup to 0.87 mgL^-1^ omethoate).

#### Pool-seq library preparation

Alive and dead mites were used to create pooled DNA extractions using a DNeasy® Blood and Tissue Kit (Qiagen, Hilden, Germany). Each DNA extraction represented a unique biological replicate comprising 10–70 mites (see Table S1). For each DNA extraction, two 150 bp paired-end whole-genome Illumina short-read libraries were constructed by Novogene (Novogene Co. Ltd, Hong Kong), which represented technical replicates of each pooled DNA extraction. Pool-seq libraries were generated for at least one pool of alive and dead mites for all populations, except Wantirna South for which we had a single untreated collection. We also produced pool-seq libraries for untreated mites from Colbinabbin Res, Colbinabbin Sus, and Manjimup, which acted as control samples.

For downstream analysis, pool-seq libraries were aggregated to different levels. For each unique DNA pool, we always aggregated replicate pool-seq libraries. In the Manjimup and Yongarillup samples, we also aggregated different DNA pools from the same population and pesticide treatment to increase the number of pooled diploid individuals. We initially extracted DNA from smaller pool sizes to increase biological replication from available mite collections. We indicate the aggregation of DNA pools from the same population and pesticide treatment in Table S1. Herein, we refer to ‘samples’ as sets of aggregated DNA pools. Samples were used to establish contrasts between alive and dead mites. All populations were represented by one alive–dead pair of samples, except the Colbinabbin Res and Sus populations, for which we had two independent pairs. For the Colbinabbin Res and Sus populations, we also included a pair of untreated samples. Details of sample contrasts are provided in Table 1.

**Table 1.** Sample contrasts for testing genetic differences between pooled *H. destructor* samples that were untreated (stochastic background differences) or treated (potential selected differences) with organophosphate.

| Sample 1 | Sample 2 | Code | Population | Contrast |
| --- | --- | --- | --- | --- |
| 2 | 3 | ColS: untreated | Colbinabbin Sus | Untreated |
| 4 | 5 | ColR: untreated | Colbinabbin Res | Untreated |
| 6 | 8 | ColS: treated 1 | Colbinabbin Sus | Treated |
| 7 | 9 | ColS: treated 2 | Colbinabbin Sus | Treated |
| 10 | 12 | ColR: treated 1 | Colbinabbin Res | Treated |
| 11 | 13 | ColR: treated 2 | Colbinabbin Res | Treated |
| 18 | 19 | Tamb: treated | Tambellup | Treated |
| 20 | 21 | Tree: treated | Treeton | Treated |
| 26\|28 | 27\|29 | Yong: treated | Yongarillup | Treated |
| 38\|39 | 40\|41 | Manj: treated | Manjimup | Treated |

**Table 2.** Single Nucleotide Polymorphisms (SNPs) within candidate *ace* target-site codons and their effects on coding potential in *H. destructor*. The canonical *ace* gene (HDE_14279) is bolded.

| Tcal | Hdes | Ace gene ID number | Contig | Position | Ref allele | Alt allele | Non-synonymous |
| --- | --- | --- | --- | --- | --- | --- | --- |
| 78 | 86 | HDE_01576 | utg155 | 546035 | V (GTA) | V (GTG) | . |
| 78 | 84 | HDE_14201 | utg9 | 345454 | V (GTG) | M (ATG) | Yes |
| 119 | 129 | HDE_09762 | utg48 | 258743 | G (GGG) | G (GGA) | . |
| **119** | **191** | **HDE_14279** | **utg94** | **20925** | **G (GGC)** | **S (AGC)** | **Yes** |
| 128 | 277 | HDE_13905 | utg83 | 1999982 | E (GAG) | E (GAA) | . |
| 150 | 159 | HDE_01556 | utg155 | 503437 | P (CCC) | P (CCA) | . |
| 201 | 210 | HDE_01553 | utg155 | 499445 | S (TCG) | S (TCT) | . |
| 201 | 207 | HDE_05340 | utg31 | 94639 | S (TCA) | S (TCG) | . |
| **201** | **272** | **HDE_14279** | **utg94** | **21168** | **A (GCC)** | **S (TCC)** | **Yes** |
| 227 | 337 | HDE_01190 | utg153 | 521028 | G (GGC) | G (GGA) | . |
| 227 | 5 | HDE_01555 | utg155 | 501653 | A (GCG) | A (GCA) | . |
| 227 | 238 | HDE_02989 | utg19 | 1187910 | G (GGA) | G (GGC) | . |
| 238 | 241 | HDE_01576 | utg155 | 546499 | R (AGG) | K (AAG) | Yes |
| 328 | 483 | HDE_13905 | utg83 | 1999364 | F (TTC) | F (TTT) | . |
| 328 | 483 | HDE_13905 | utg83 | 1999366 | M (ATG) | V (GTG) | Yes |
| 331 | 97 | HDE_01555 | utg155 | 501929 | L (CTC) | L (CTG) | . |
| 331 | 326 | HDE_01576 | utg155 | 546755 | H (CAC) | H (CAT) | . |
| **331** | **401** | **HDE_14279** | **utg94** | **21556** | **F (TTC)** | **Y (TAC)** | **Yes** |
| 336 | 450 | HDE_01190 | utg153 | 521367 | F (TTT) | F (TTC) | . |
| **336** | **406** | **HDE_14279** | **utg94** | **21572** | **L (CTA)** | **L (CTG)** | . |
| 396 | 544 | HDE_07881 | utg40 | 396829 | G (GGC) | G (GGT) | . |
| 441 | 589 | HDE_07881 | utg40 | 396694 | G (GGC) | G (GGA) | . |

Note: Codons are reported with respect to their candidate *ace* position in *T. californica* ('Tcal') and their relative *H. destructor* codon ('Hdes'). Alleles are reported with respect to their amino acid residue, with nucleotide bases comprising the codon in brackets.

#### Bioinformatic processing and SNP variant calling

Raw reads for each unique pool-seq library were quality trimmed using FASTP (Chen, Zhou, Chen, & Gu, 2018), truncating reads to 100 bp and discarding all reads below this threshold. Trimmed reads for each pool-seq library were mapped to the *H. destructor* reference genome (GenBank GCA_022750525.1) using BOWTIE2 (Langmead & Salzberg, 2012) with a global alignment and the ‘--sensitive’ mapping option. Following trimming, our FASTQ files for individual pool-seq libraries contained a mean of 29,570,401 reads with a range of 21,791,964–41,049,660 reads. The mean number of reads mapping to the*H. destructor* genome (genomic contigs >0.2 Mb) was 15,522,450 reads with a range of 11,132,739–23,519,859 per individual pool-seq library. Mapping was limited to contigs >0.2 Mb in length because our prior study identified that mapping to contigs <0.2 Mb led to variable and unreliable mapping (Thia et al., 2023). SAMTOOLS (Danecek et al., 2021; H. Li et al., 2009) was used to discard reads with mapping quality <30 and those that were found to be PCR duplicates. Variant calling was performed using FREEBAYES (Garrison & Marth, 2012) with the arguments: ‘--min-alternate-count 3’ and ‘--pooled-continuous’. The FREEBAYES script, *fasta_generate_regions.py*, was used to segment the genome for variant calling in parallel. We used BCFTOOLS (Danecek et al., 2021; H. Li, 2011) to combine VCF files from across the parallel variant calling runs. Variant filtering was performed with VCFTOOLS (Danecek et al., 2011) to retain only single nucleotide polymorphisms (SNPs; no indels) with a minor allele count of 3, a minor allele frequency of 0.01, minimum quality of 30, minimum depth of 20 reads per sample, no missing data, and to exclude SNPs from repetitive regions of the genome.

#### Copy number variation

We estimated a proxy for copy number at the canonical *ace* and *ace*-like genes using whole-genome data. These estimates of copy number were then used to test the hypothesis that greater copy number is associated with organophosphate insensitivity in *H. destructor*. Our proxy for copy number at the canonical *ace* or *ace*-like genes was derived from read depth ratios relative to 866 single copy reference genes that were verified in prior BUSCO analyses (Thia et al., 2023). These single copy reference genes all resided on genomic contigs >0.2 Mb in length. We extracted mean read depth information for these single copy reference genes, the canonical *ace*, and 18 *ace*-like genes with the SAMTOOLS COVERAGE function. Read depth statistics were then imported into R v4.1.3 (R Core Team, 2022) for further analysis.

We first summarized read depth for the canonical *ace*, the 18 focal *ace*-like genes, and the 866 single copy reference genes as the mean across replicates libraries nested in pools nested in samples (Table S1). We then obtained read depth ratios in the canonical *ace* or *ace*-like genes as a proxy for copy number by dividing their read depth by that of each single copy reference gene. The *ace* and *ace*-like genes were distributed across 11 genomic contigs, and the single copy reference genes were distributed across 52 genomic contigs. All genomic contigs containing the focal canonical *ace* and *ace*-like genes were represented in contigs containing single copy reference genes. This meant we were able to incorporate genome-wide variation in read depth into our formal hypothesis using a mixed effects linear model:

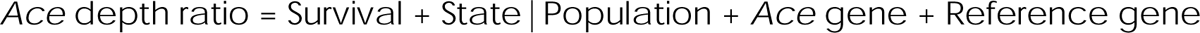

In this model, the response ‘*Ace* depth ratio’ is the read depth ratio for the canonical *ace* or *ace*-like genes. ‘Survival’ is the survival status, a categorical fixed effect, alive or dead.We fitted the random intercepts for ‘Population’ nested within ‘State’ of sample origin, and separate random intercepts for the canonical *ace* or *ace*-like genes, ‘*Ace* gene’, and each single copy reference gene, ‘Reference gene’. The model was fitted using the LME4 (Bates, Mächler, Bolker, & Walker, 2014) and LMERTEST (Kuznetsova, Brockhoff, & Christensen, 2017) R packages. We calculated the mean difference in depth ratio across *ace* genes to summarise differences in depth ratio between alive and dead mites.

### Target-site mutations

We surveyed the canonical *ace* and *ace*-like genes for polymorphisms at codons known to be associated with target-site resistance in other arthropods: 73, 78, 82, 119, 128, 129, 150, 201, 201, 227, 280, 290, 328, 330, 331, 336, 435, and 441*T*(*orpe do californica* numbering) (reviewed in: Fournier, 2005; Feyereisen*et al*., 2015). We used our alignment of *H. destructor ace* protein sequences with *T. californica* to determine the homologous codon positions in *H. destructor*. We then identified the underlying nucleotide bases for each codon and determined whether any of our SNPs were found within these codons. We surveyed the 19 *ace* genes residing on genomic contigs >0.2 Mb in length. To further explore the distribution of amino acid polymorphisms, we examined the read alignments in a subset of pool-seq libraries. We extracted mapped reads from select libraries from Wantirna South, Colbinabbin Sus, Colbinabbin Res, and Manjimup populations using the G ENOMI C ALI GNMENTS (Lawrence et al., 2013) and BIOSTRINGS (Pagès, Aboyoun, Gentleman, & DebRoy, 2017) R packages.

After identifying non-synonymous SNPs in the a priori target-site codons, we tested whether polymorphisms at these SNP were associated with organophosphate insensitivity. We compared the allele frequencies between alive and dead mites using a Χ^2^ test of the observed read counts for the reference and alternate allele (a two-by-two contingency table). We calculated the frequency differential between treated alive and dead mites (or between untreated samples) in each sample pair (Table 1) by subtracting the reference allele frequency of dead mites from that of alive mites (or one untreated control sample from another).

#### Genomic outlier scan

We used our whole-genome data to perform a scan for outlying genomic regions that may exhibit signatures of organophosphate selection when comparing alive and dead mites. The purpose of this analysis was two-fold: (1) to provide additional support for the contributions of the canonical *ace* or other *ace*-like genes to organophosphate insensitivity; and (2) to identify other non-*ace* genes that may contribute to organophosphate insensitivity. We measured genetic differentiation between alive and dead mites using the mean *F*_ST_ for a genomic region (‘blocks’). We aggregated pool-seq libraries to generate reference and alternate allele read counts for working sample pairs in R (Table 1). We used the POOLFSTATR package to filter SNPs. For each working sample pair, SNPs were filtered at a minor allele frequency threshold of 0.1. We then defined SNP blocks as genomic regions comprising 30–50 SNPs, resulting in a median SNP block size of 6,040 bp, and a range of 1,022–82,863 bp. We also performed these same tests on untreated sample pairs to infer the stochastic background differentiation expected.

To calculate the mean *F*_ST_ for a genomic region, we first calculated SNP-wise estimates of *F*_ST_ ‘corrected’ for sample size and technical error using POOLFSTAT:

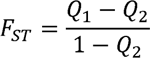

Here, *Q*_1_ is the identity in state probability for two genes sampled within a population pool, and *Q*_2_ is the identity in state probability for two genes sampled between two population pools. This formulation is equivalent to θ, as defined by Weir & Cockerham’s (1984). We also calculated an ‘uncorrected’ *F*_ST_ (Weir & Cockerham, 1984):

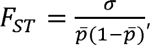

where σ is the variance in the reference allele frequency and is the mean reference allele frequency. After calculating SNP-wise estimates of *F*_ST_, we calculated block-wise *F*_ST_ by dividing the mean numerator by the mean denominator variance components for all SNPs within a block. We therefore had block-wise *F*_ST_ estimates corrected and uncorre cted for sample size and technical error. SNP blocks exhibiting unusually high genetic differentiation were identified using a two-step method. The uncorrected block-wise*F*_ST_ values were subjected to the algorithm described in the OUTFLANK method (Whitlock & Lotterhos, 2015) to identify a set of ‘preliminary’ outlier blocks. To execute the OUTFLANK method, we used a hybrid script comprised of custom code and functions borrowed from the O UTFLANK GitHub (used to calculate effective degrees of freedom and the two-sided Χ^2^ test-statistic *p* value) (Whitlock, 2019). The preliminary outlier blocks were retained if they exceeded the 99^th^ percentile calculated from corrected block-wise *F*_ST_ values. Those blocks passing both steps comprised our ‘final’ outlier blocks.

Genes residing in the final outlier blocks were considered candidate genes of organophosphate insensitivity. To test whether certain types of genes were over-represented in this candidate gene set, we used a gene ontology (GO) enrichment analysis. GO enrichment was analyzed using Fisher’s exact tests in R. The significance of enrichment was assessed following Bonferroni correction of *p* values.

## Results

### Copy number variation

We found support for our hypothesis that greater copy numbers of the canonical *ace* and *ace*-like are associated with organophosphate insensitivity, but the role of copy number may vary by gene and (or) population. Most genes had read depth ratios that were distributed around 1.00, suggesting they existed largely as single copies, but a few genes had depth ratios that deviated notably from 1.00 in some or all samples (Figure 3). In the canonical *ace*, HDE_14279, samples from three Western Australia populations, Manjimup, Treeton, and Yongarillup had depth ratios ranging from 1.49–2.37, suggesting a much higher average copy number in these populations. In contrast, several *ace*-like genes had depth ratios <1.00 across all samples, suggesting that these genes might not be present in every genome: HDE_12794 (0.55–0.86), HDE_09715 (0.16–0.50), HDE_09713(0.36–0.64), HDE_09712 (0.53–0.82), and HDE_09711 (0.73–0.91).

**Figure 3.**
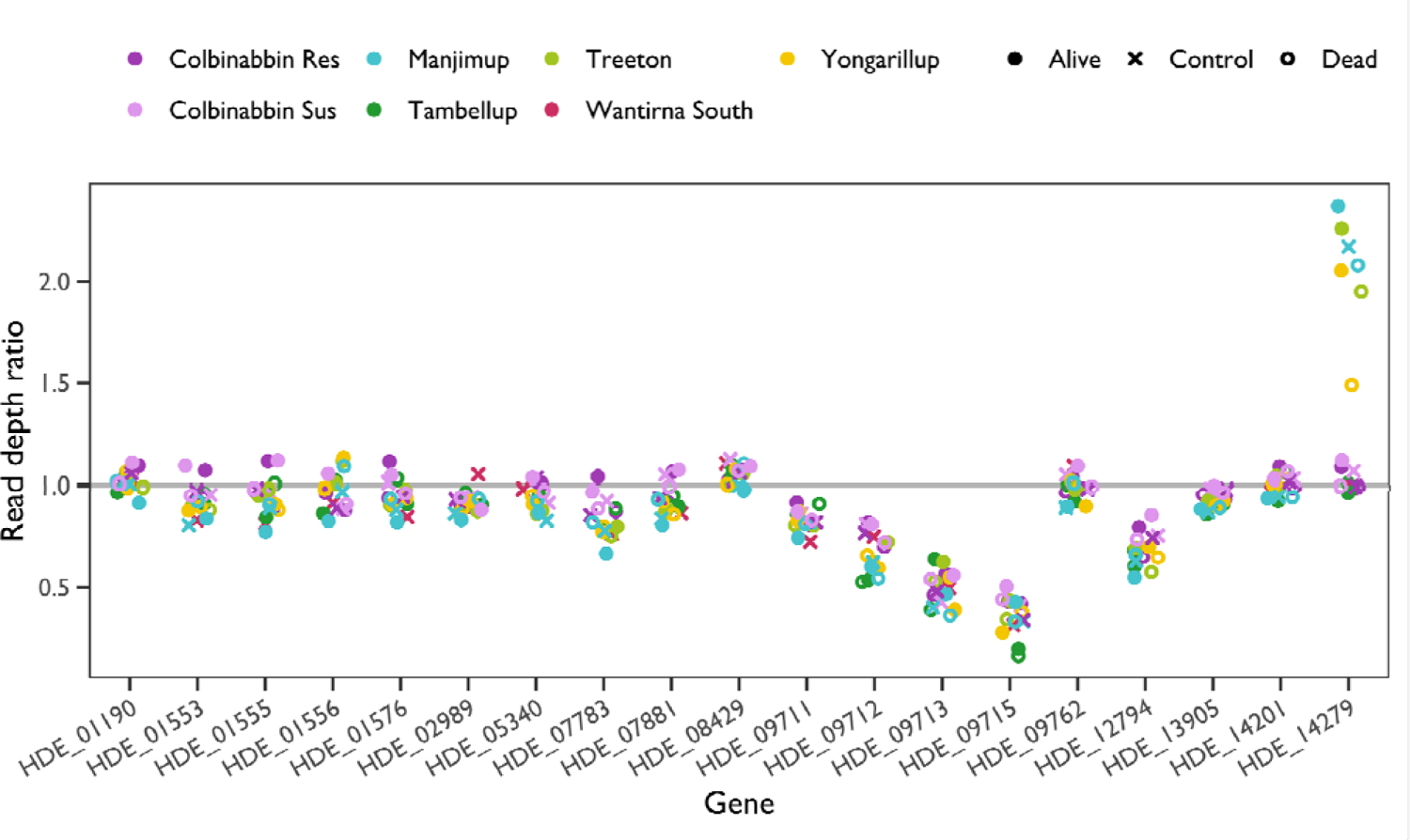
Mean copy number at the canonical ace and *ace*-like genes in population pool seq samples of *H. destructor*. The *x*-axis represents the 18 focal ace-like genes and the canonical *ace* (HDE_14279). The *y*-axis represents the mean read depth ratio (a proxy for copy number), which is the read depth of the canonical or *ace*-like gene divided by the read depth of 866 single copy reference genes. Points represent samples: colors and shapees denote population and survival status (see legend). The grey horizontal line demarcates a read depth ratio (copy number) of one.

Our linear mixed effects model of read depth ratio identified a significant fixed ‘Survival’ effect (*F*_1,196558_= 749.13, *p* < 0.001), but the overall effect was small (β_Dead_ = –0.02, *t* = –27.37, df = 196558, *p* < 0.001). On average, alive mites had a read depth ratio of 0.91 (±0.06 SE), whereas dead mites on average had a read depth ratio of 0.89 (± 0.0007 SE). We also reran the model excluding the canonical *ace* gene to check that it was not biasing results. There was still an overall positive association between copy number in *ace*-like genes and survival following organophosphate exposure (*F*_1,186167_= 390.46, *p* < 0.001), although the effect size was smaller after removing the canonical *ace* (β_Dead_ = –0.007, *t* = –19.76, df = 1861670,*p* < 0.001). The mean difference in depth ratios between alive–dead pairs were positive for the Colbinabbin Res (0.09 ± 0.01 SE), Colbinabbin Sus (0.07 ± 0.01 SE), Treeton (0.04 ± 0.02 SE), and Yongarillup (0.09 ± 0.01 SE) populations. In contrast, the mean difference in depth ratios was negative for the Manjimup (–0.05 ± 0.03 SE) and Tambellup (–0.05 ± 0.02 SE) populations.

These results suggest that at least some *ace*-like genes may have subtle contributions to organophosphate insensitivity through small variations in copy number (Figure 4). In a subset of populations, a more discernible increase in copy number in the canonical *ace* gene is linked to organophosphate insensitivity, which may help over-express resistant proteins (Figure 4; also see below).

**Figure 4.**
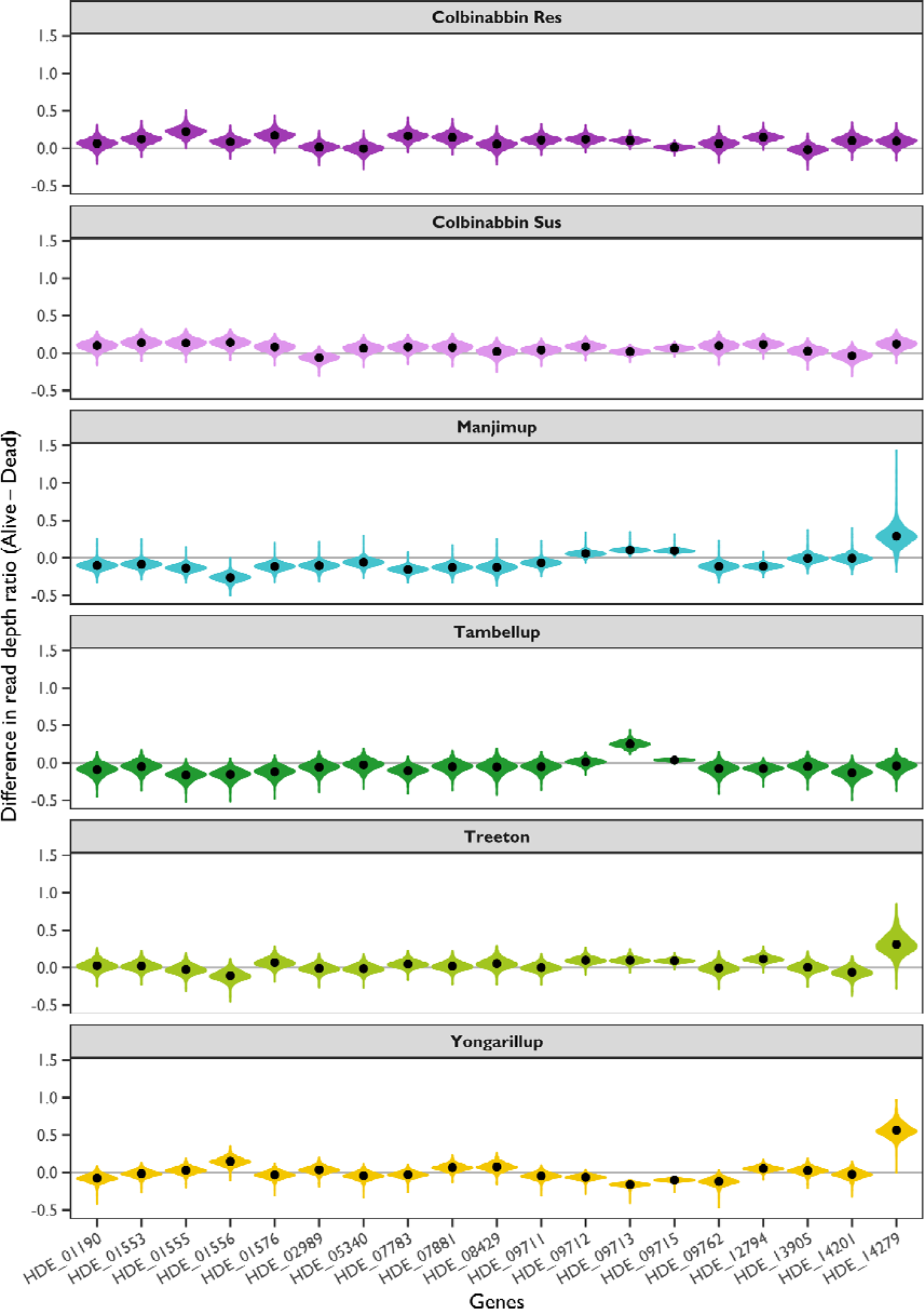
Differences in canonical *ace* or *ace*-like gene copy number between alive and death mites following organophosphate exposure. The*x*-axis represents the 18 focal ace-like genes and the canonical *ace* (HDE_14279). The *y*-axis represents the difference in gene copy number as estimated by the read depth ratio between alive and dead mites (alive – dead). Black points represent the mean difference relative to estimates from 866 single copy reference genes. Violin plots summarise the distribution of estimates. The grey line demarcates a difference of zero. Each panel represents a population.

### Target-site mutations

For the 19 target-sites codons surveyed, only 6 non-synonymous SNPs were identified across all samples (Table 2). One SNP was observed in *ace*-like HDE_01576, a R238K mutation; one SNP was observed in *ace*-like HDE_13905, a M328V mutation; one SNP was observed in*ace*-like HDE_14201, a V78M; and three SNPs were observed in the canonical *ace* HDE_14279, producing G119S, A201S, and F331Y mutations. Despite the minimal within-gene level polymorphism, there was considerable among-gene level polymorphism (Figure 5). Across the 19 analyzed genes, there was an average of 7.1 variants (amino acids or deletions) at target-site codons, with a range of 3 to 13 variants. Particularly interesting was the observation that some *ace*-like genes in *H. destructor* were fixed for amino acids that are known to confer resistance in other arthropods (Feyereisen et al., 2015). Most striking were the following codons: codon 119, with 6/17 genes fixed for either A119 or S119 (Figure S2); codon 201, with 12/17 genes fixed for S201 (Figure S3); codon 227, with 8/18 genes fixed for A227 (Figure S4); codon 331, with 2/18 genes fixed for H331 (Figure S5); and codon 441, with 6/18 genes fixed for G441 (Figure S6). These fixed alleles that are known to confer resistance were found in both susceptible and resistant populations. Note that in some of the analyzed genes an equivalent target-site codon was absent, hence *n* ≠ 19.

**Figure 5.**
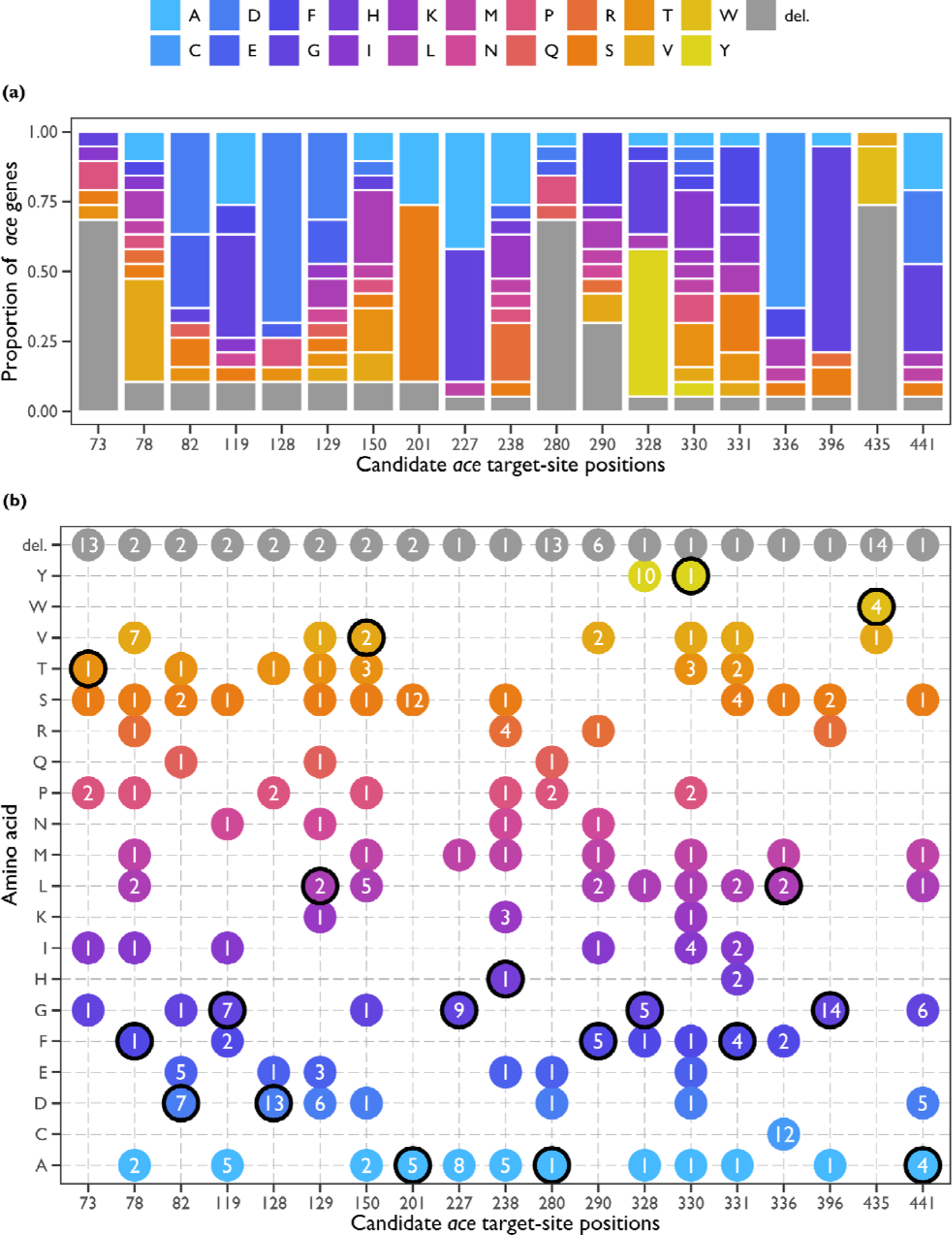
Reference amino acids across the *H. destructor* canonical *ace* and focal 18 *ace* like genes (on contigs >0.2 Mb) at candidate target-site codons (*T. californica* numbering). (a) Amino acid proportions. The *x*-axis is the candidate *ace* target-site codon, the *y*-axis is the relative proportion of *ace* genes exhibiting a particular amino acid as their reference allele (see legend) in the *H. destructor* genome. (b) Amino acid counts. The *x*-axis is the candidate *ace* target-site codon, the *y*-axis is the *ace* gene numbers exhibiting a particular amino acid as their reference allele in the *H. destructor* genome. A black outline denotes the amino acid present in the canonical *ace* gene, HDE_14279. (a, b) ‘del.’ indicates a deletion, that is, where there is no equivalent codon in the *H. destructor* gene relative to *T. californica*.

We found mixed support for the hypothesis that target-site mutations are associated with organophosphate insensitivity. If a mutation conferred insensitivity, we would expect that the difference in mutant allele frequency (alive minus dead) would be positive. The non-synonymous mutations at the *ace*-like genes have not previously been associated with organophosphate resistance and were found in both resistant and susceptible populations. However, the G119S, A201S, and F331 mutations at canonical *ace* HDE_14279 are known resistance-conferring mutations and were only observed in the resistant populations in our study, although their direction of effect was not consistent (Figure 6a).

**Figure 6.**
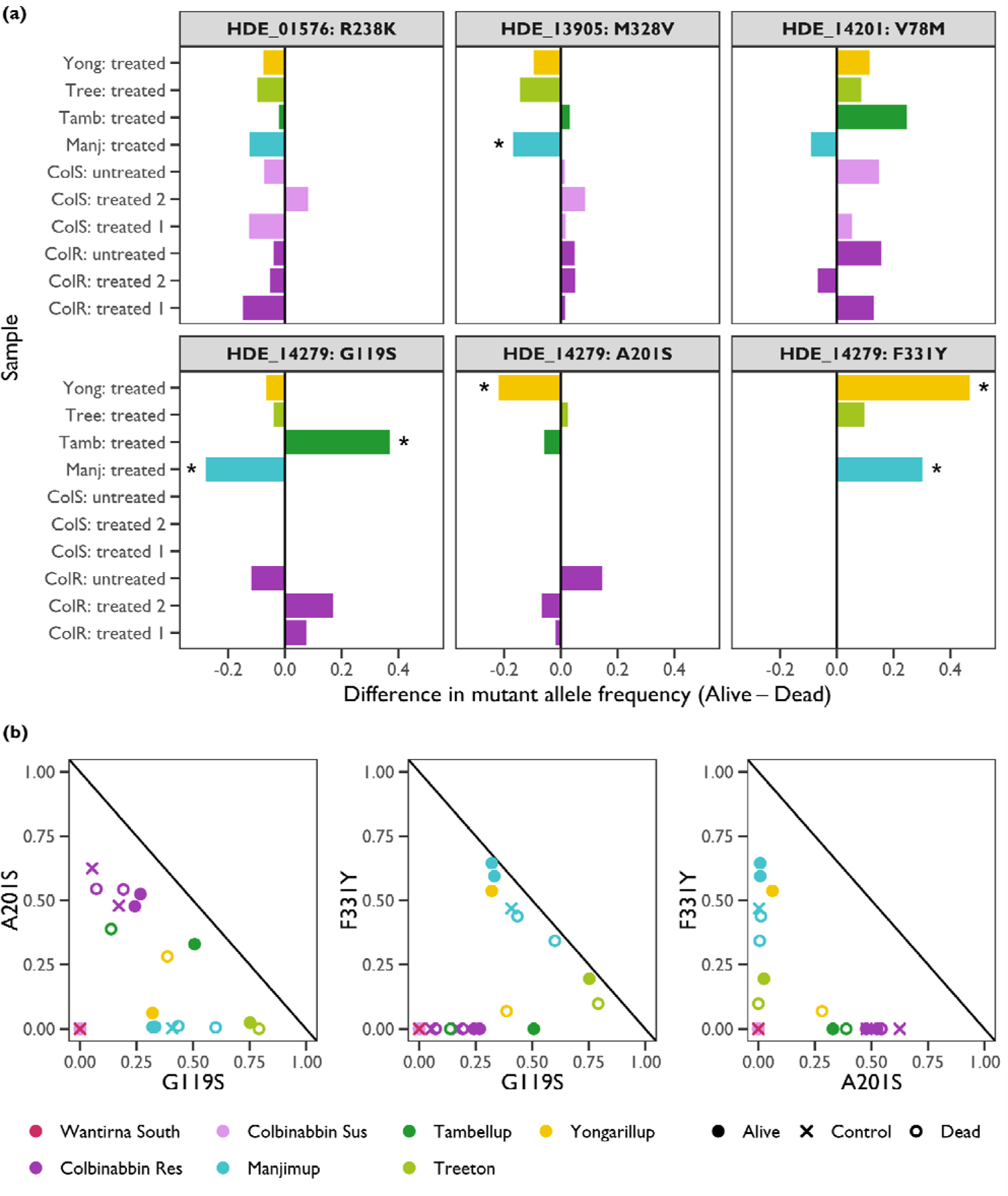
Allelic variation for non-synonymous mutations at candidate *ace* target-sites cod ns in *H. destructor* (with *Torpedo californica* numbering). (a) Mutant allele frequency differenc e between alive and dead exposed to organophosphates (‘treated’) or between two unexposed samples (‘untreated’). Bars on the *x*-axis represents the allele frequency difference and populations are organized on the *y*-axis. Positive values on the *x*-axis indicate samples of alive mites had a higher mutation allele frequency relative to samples of dead mites. Asterisks denote significant allele count differences following Bonferroni correction for multiple tests (χ^2^ test, α = 0.05/50 = 0.001). Population codes are: ‘ColR’ = Colbinabbin Res, ‘ColS’ = Colbinabbin Sus, ‘Manj’ = Manjimup, ‘Tamb’ = Tambellup, ‘Tree’ = Treeton, and ‘Yong’ = Yongarillup. Each panel represents a gene and mutation; note that the canonical *ace* gene is HDE_14279. (b) Mutant allele frequency correlations for target-site mutations at the canonical *ace* gene, HDE_14279. The *x*-axis and *y*-axis represent the frequencies of one of three target-site mutations. The 1:1 negative correlation is depicted by the black line. Symbols represent samples, with colours denoting population, and shapes denoting survival group (see legend).

The mixed patterns of differentiation in G119, A201S, and F331Y mutations in canonical *ace* are probably due to different haplotypes being selected by organophosphates. This was evidenced by negative correlations in the frequencies of target-site mutations (vs. the expected positive correlation if these mutations were linked) (Figure 6b). The A201S and F331Y mutations did not occur (or only rarely occurred) in the same resistant population: populations that were polymorphic for one of these mutant alleles were fixed (or almost fixed) for the wild-type allele at the other site (Figure 6b, right panel). In contrast, the G119S mutation was negatively correlated with A201S and (or) F331Y in some populations where there was polymorphism for these mutations. For Colbinabbin Res, Tambellup, and Yongarillup, as the G119S mutant allele frequency increased, the A201S mutant allele frequency decreased (Figure 6b, left panel). For Manjimup, Treeton, and Yongarillup, as the G119S mutant allele frequency increased, the F331Y mutant allele frequency decreased (Figure 6b, middle panel). The apparent observable phases for resistant mutations based on allele frequencies appear to be: G119/S201, S119/A201, G119/Y331, S119/F331. However, the higher copy number at the canonical *ace* in Manjimup, Treeton, and Yongarillup complicates the picture; there may be haplotypes with different copy numbers segregating with different mutations in copies.

### Genomic outlier scan

Two key results emerged from our genomic outlier scan: (1) signatures of selection were not discernible at the canonical *ace* or *ace*-like genes; and (2) there was no evidence of other enriched functions involved in organophosphate insensitivity. On average, corre cted block-wise *F*_ST_ between alive–dead sample pairs ranged from 0.008 to 0.035, with a mean of 0.0192, across the different populations (Figure S7). For Colbinabbin Res and Colbinabbin Sus, we had two independent untreated sample pairs that allowed us to assess the expected background corrected block-wise *F*_ST_, which was 0.012 for both populations. Genetic differentiation between untreated samples was larger or equivalent to treated alive–dead sample pairs in both the Colbinabbin Res and Colbinabbin Sus populations. Any genetic differentiation resulting from organophosphate selection in our experiments are unlikely to be detected above background stochastic sampling noise in allele frequencies.

Across our samples pairs used in genomic outlier scans, in only one instance was an *ace*-like gene recovered (HDE_01553), but this was for an untreated sample pair (Colbinabbin Sus, sample 5 vs 4) where neither sample had been exposed to organophosphates. The genomic block containing the canonical *ace* was never significantly differentiated. A total of 474 unique genes were identified in outlier genomic blocks, but only 17 were shared by two populations. There was very little evidence for significant GO term enrichment in any alive– dead sample pair after accounting for multiple testing. Of those significant GO terms, ‘Mannose metabolic process’ and ‘alpha-mannosidase activity’ were enriched in the Colbinabbin Res ‘treated 2’ sample pair, and ‘microtubule-based process’ and ‘dyein complex’ were enriched in the Manjimup ‘treated’ sample pair. None of these GO terms have clear links to pesticide resistance.

## Discussion

Our study suggests that a combination of copy number variation and target-site mutations may contribute to organophosphate insensitivity in Australian populations of *H. destructor*. Increased copy number can provide pesticide insensitivity through dosage effects (Kwon, Clarkt, & Lee, 2010; Deok Ho Kwon, Choi, Je, & Lee, 2012). At the canonical *ace* gene, we observed evidence of high mean copy number (≥2) in a subset of populations from Western Australia, which may help over-express target-site mutations (G119S, A201S, and F331Y) that are segregating at this gene. Such a mechanism has been observed in other arthropod pests, such as *Tetranychus evansi*, the tomato red spider mite (Carvalho et al., 2012), and *Aphis gossyppii*, the cotton aphid (Shang et al., 2014), and it is also common in organophosphate-resistant mosquito species (Weetman, Djogbenou, & Lucas, 2018).

Whilst the target-site mutations at the canonical *ace* are only segregating in resistant populations of *H. destructor*, they did not show clear patterns following organophosphate exposure. Indeed, even mites that were killed by organophosphates exhibited relatively high frequencies of G119S, A201S or F331Y at canonical *ace*. We were also unable to detect any signatures of selection at canonical *ace* from our genomic outlier scans. Multiple factors might have reduced our capacity to observe a clear signal in the canonical *ace* gene. First, the effect of the target-site mutations may not be large, which contrasts the pyrethroid resistance mechanism in *H. destructor* (Edwards et al., 2018; Yang et al., 2020), and the clear effects of these same *ace* target-site mutations in other arthropod pests (Andrews, Callaghan, Field, Williamson, & Moores, 2004; Carvalho et al., 2012; Khajehali et al., 2010). Second, target-site mutations could vary in their effects across different genetic backgrounds (Li et al., 2015; Riga et al., 2017). Third, haplotypes with different combinations of copy number and target-site mutations may vary in their effect sizes, such that organophosphates are not simply selecting for alleles at the canonical *ace*, but for different haplotypes. Long-read sequencing would be beneficial for resolving which haplotypes respond most to selection in *H. destructor*. We also suspect that the variation in the canonical *ace* is only one contributing component of a polygenic architecture of organophosphate insensitivity. Further genomic investigations are required to discern the possible roles of detoxification (*e.g.,* cytochrome P450s, GSTs, esterases, or ABC transporters) or reduced penetrance (*e.g.,* cuticular proteins) mechanisms to organophosphate insensitivity in *H. destructor*.

The radiated *ace*-like genes may have more subtle contributions to organophosphate insensitivity. We have shown previously that these *ace*-like genes may be involved in complex haplotypes and structural variants, which may result in the presence or absence of different genes among individuals (Thia et al., 2023). In this study, we provide further support for this hypothesis by demonstrating apparent copy numbers <1 for some *ace*-like genes. We also observed a significant association between copy number and organophosphate insensitivity, which suggests that surviving mites had slightly higher copies of (some) radiated *ace*-like genes. Amino acids at target-site positions were variable among *ace*-like genes. Duplication may relax evolutionary constraints on genes, allowing them to evolve more freely and accumulate mutations (Lynch & Conery, 2000). However, in some cases, we noticed a high proportion of known resistance-conferring mutations at these positions. The most striking was position 201, a major residue at the *ace* catalytic site, which forms part of the oxyanion hole (Oakeshott et al., 2005). At this site, the only alleles were either A201 (wild-type) or S201 (mutant), for the canonical *ace* and all *ace*-like genes analysed; this is intriguing given the diversity observed at other sites and the known role of S201 in resistance. We note that the contribution of *ace*-like genes to organophosphate insensitivity are likely subtle: (i) estimated coefficients from our linear models were small; (ii) *ace*-like genes were not recovered from our genomic outlier scans; and (iii) radiated *ace*-like genes are found in organophosphate resistant and susceptible populations. We are currently exploring functional analyses to better understand how *ace*-like genes might contribute to organophosphate insensitivity through possible sequestration or breakdown of pesticide molecules (Kim et al., 2012; Newcomb et al., 1997).

Given the accelerating spread of organophosphate resistance across Australian populations of *H. destructor* (Arthur et al., 2021), clear diagnostic genetic markers of resistance would be highly advantageous (Thia, Hoffmann, & Umina, 2021). At present, the canonical *ace* gene remains the strongest single candidate for developing molecular diagnostic of resistance risk in Australian populations of *H. destructor*. Our work points to variable ways in which the canonical *ace* and other *ace*-like genes in *H. destructor* may contribute to organophosphate insensitivity. The combination of canonical *ace* and radiated *ace*-like genes with other non-*ace* genes may allow populations of *H. destructor* to evolve diverse responses to organophosphate pesticides. The potential non-convergence and polygenicity of adaptation to organophosphates in *H. destructor* will need to be considered when using diagnostics of resistance in molecular monitoring programs.

## Acknowledgements

This work was funded by the Grains Research Development Corporation (Australia) as part of the Australian Grains Pest Innovation Program, and with support from the University of Melbourne. Computational resources were provided by The University of Melbourne’s Research Computing Services and the Petascale Campus Initiative. We thank our three anonymous reviewers for their comments, which greatly improved this manuscript.

## Data accessibility

All sequencing data has been submitted under the GenBank Project PRJNA756307 and their accessions are listed in Table S1. All code and data required to run these analyses have been deposited into a Dryad archive (Thia, 2023): https://doi.org/10.5061/dryad.dr7sqvb2m

## Author contributions

JT formulated the original project design. PU and AH contributed to project design and obtained funding. JT performed all analyses. JT wrote the original manuscript. PU and AH contributed toward revisions of the final manuscript.

## Supplementary information

**Figure S1.**
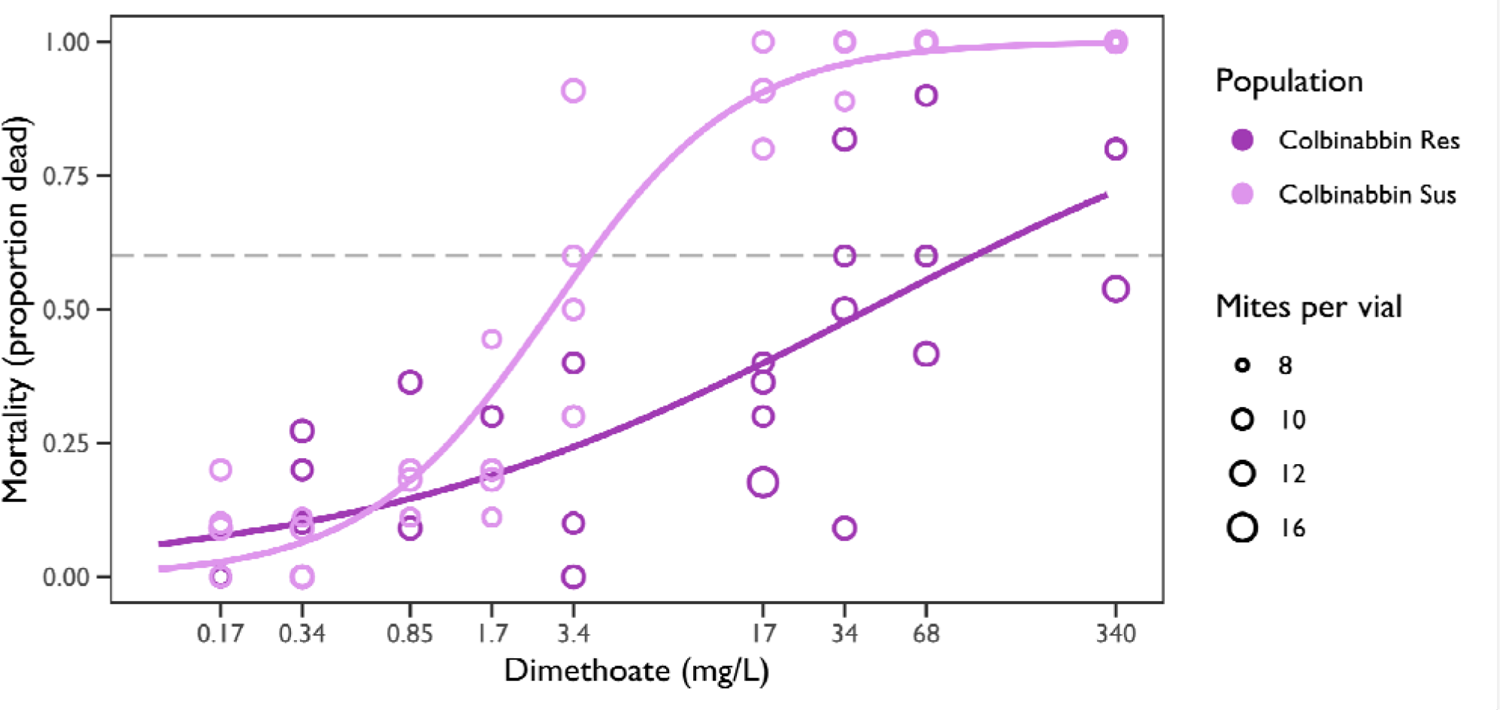
Dose-responses of the Colbinabbin populations of *Halotydeus destructor* to dimethoate. Data were from preliminary assays to establish LD60 concentrations for artifici l selection. Each population was assayed at 10 doses of dimethoate using the coated vial method. Each dose was replicated by four vials, with a mean of 10 mites per vial and a ra ge of 8–17 mites. Mites were stored at 11°C and scored after 24 hours. The *x*-axis is the dose of dimethoate and the *y*-axis is the proportion of mortality. Points represent replicate vials, with point size depicting the number of mites per vial (see legend). Solid curves represent the estimated dose-response relationship estimated from logistic regressions fitting mortality asa function of log10-transformed dose with a quasi-binomial error distribution. Points and curv s are coloured by population (see legend). The grey dashed horizontal line depicts the LD60 threshold.

**Figure S2.**
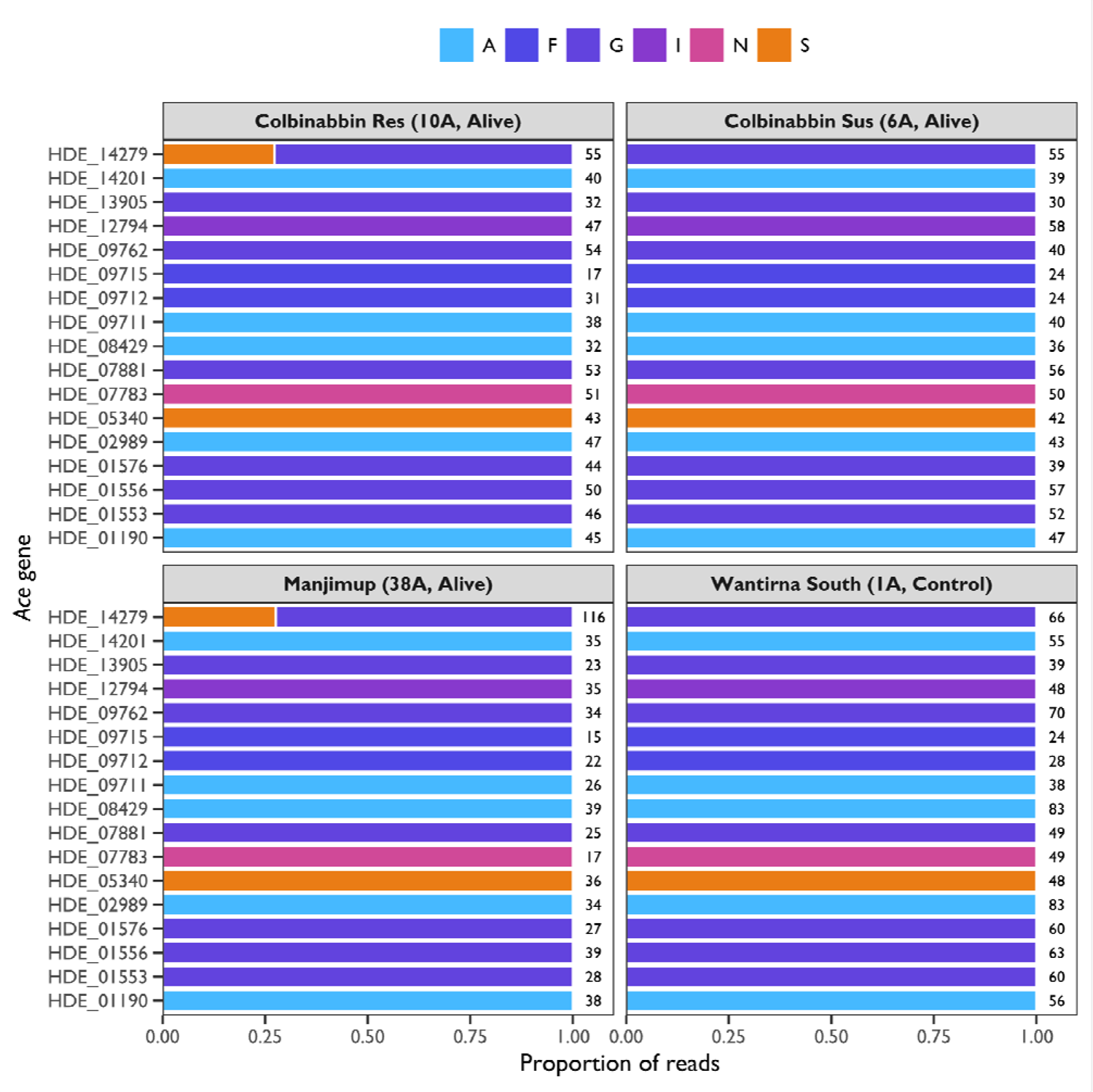
Proportion of reads for different amino acids at codon 119 (*Torpedo californica* numbering) across *ace* genes in *Halotydeus destructor*. The *x*-axis is the proportion of reads and the *y*-axis the *ace* gene. Horizontal bars represent the proportion of reads with nucleotide sequences encoding different amino acids, see legend. Numbers to the right of the bars indicate the number of observed reads that completely cover ed the codon. Paneels contain observations for select pool-seq libraries, read as ‘Population (DNA pool, Survival group)’. Note, Colbinabbin Res and Manjimup are resistant to organophosphates, whereas Colbinabbin Sus and Wantirna South are susceptible.

**Figure S3.**
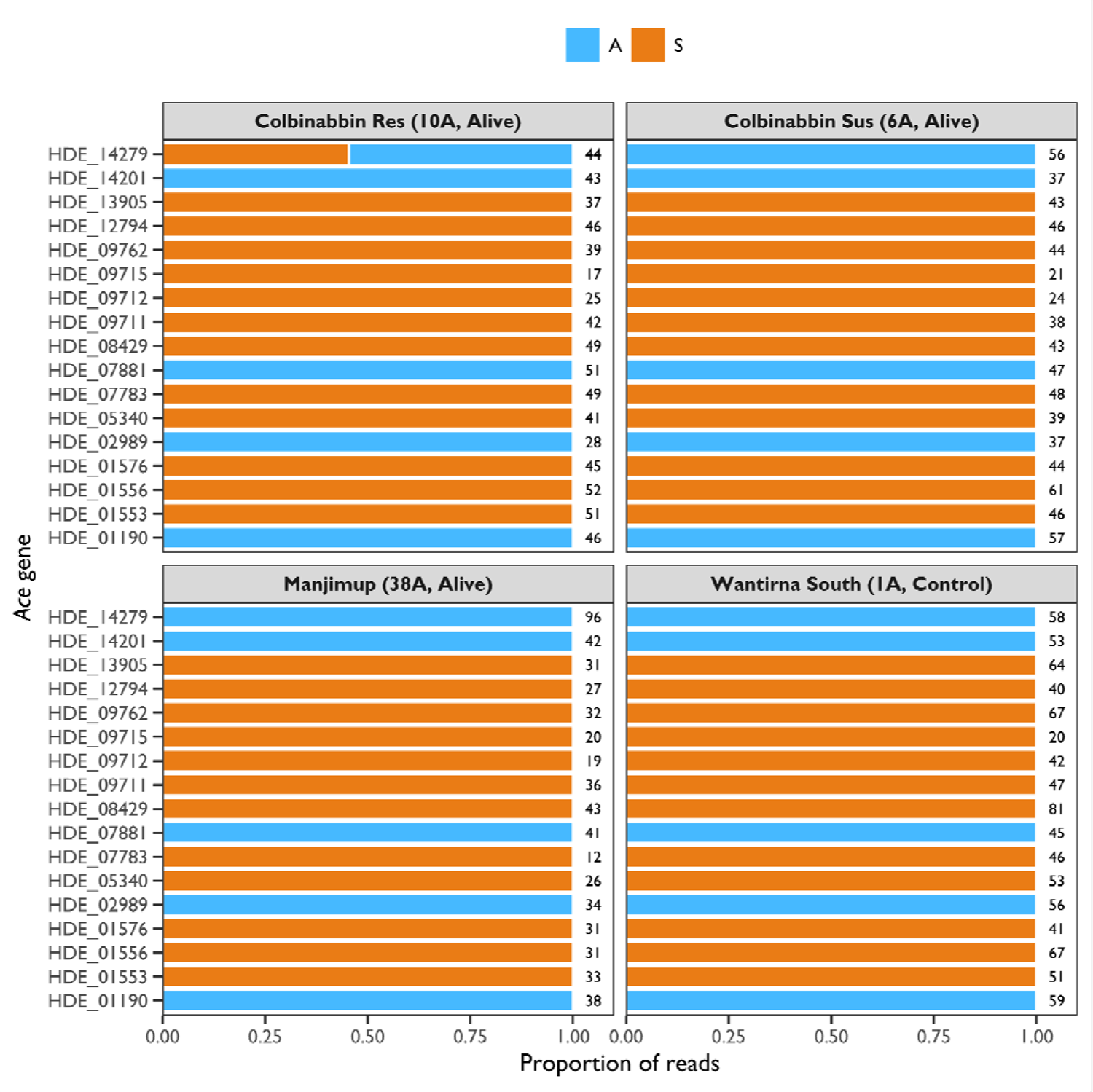
Proportion of reads for different amino acids at codon 201 (*Torpedo californica* numbering) across *ace* genes in *Halotydeus destructor*. The *x*-axis is the proportion of reads and the *y*-axis the *ace* gene. Horizontal bars represent the proportion of reads with nucleotide sequences encoding different amino acids, see legend. Numbers to the right of the bars indicate the number of observed reads that completely covered the codon. Paneels contain observations for select pool-seq libraries, read as ‘Population (DNA pool, Survival group)’. Note, Colbinabbin Res and Manjimup are resistant to organophosphates, whereas Colbinabbin Sus and Wantirna South are susceptible.

**Figure S4.**
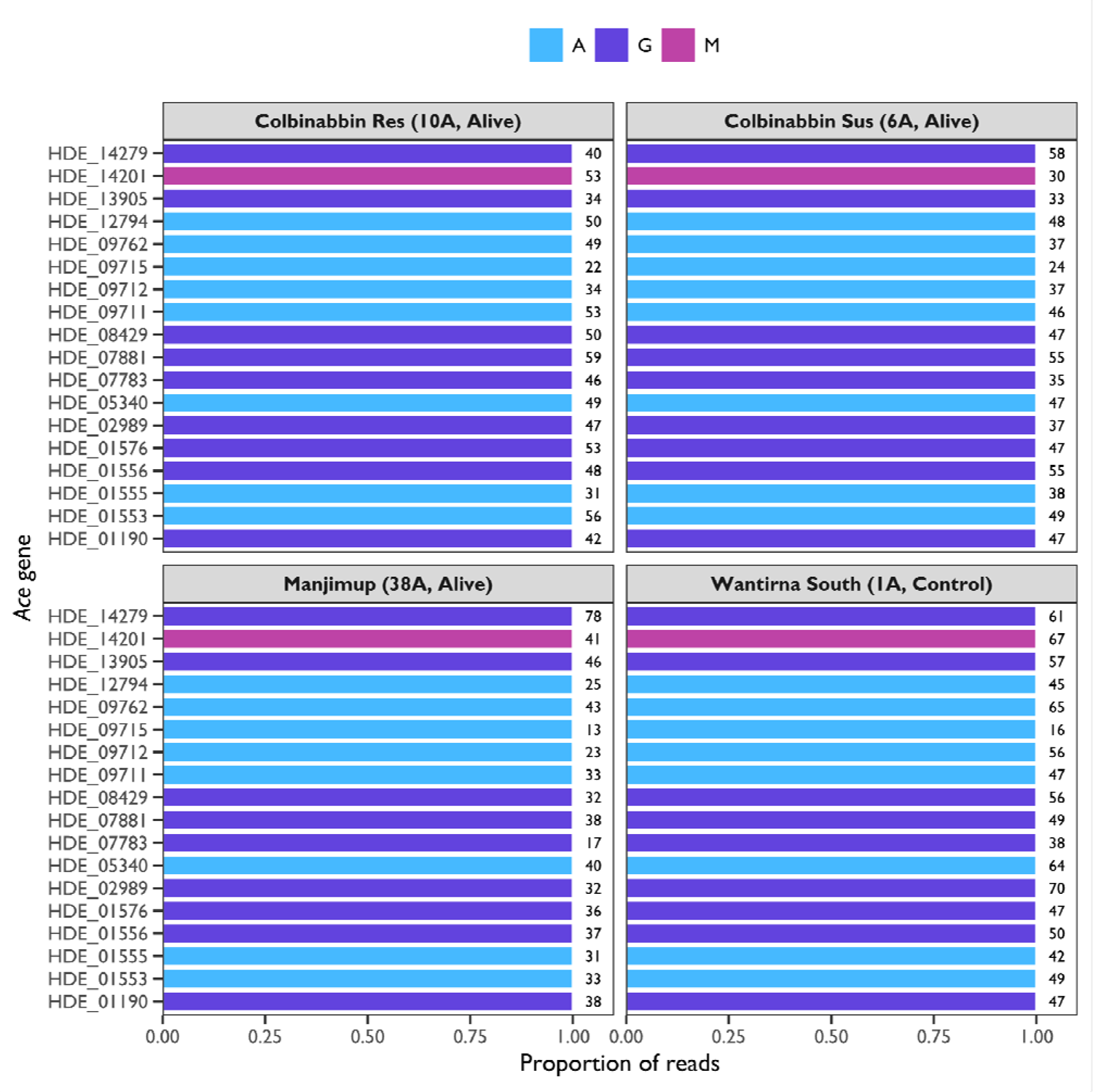
Proportion of reads for different amino acids at codon 227 (*Torpedo californica* numbering) across *ace* genes in *Halotydeus destructor*. The *x*-axis is the proportion of reads and the *y*-axis the *ace* gene. Horizontal bars represent the proportion of reads with nucleotide sequences encoding different amino acids, see legend. Numbers to the right of the bars indicate the number of observed reads that completely covered the codon. Paneels contain observations for select pool-seq libraries, read as ‘Population (DNA pool, Survival group)’. Note, Colbinabbin Res and Manjimup are resistant to organophosphates, whereas Colbinabbin Sus and Wantirna South are susceptible.

**Figure S5.**
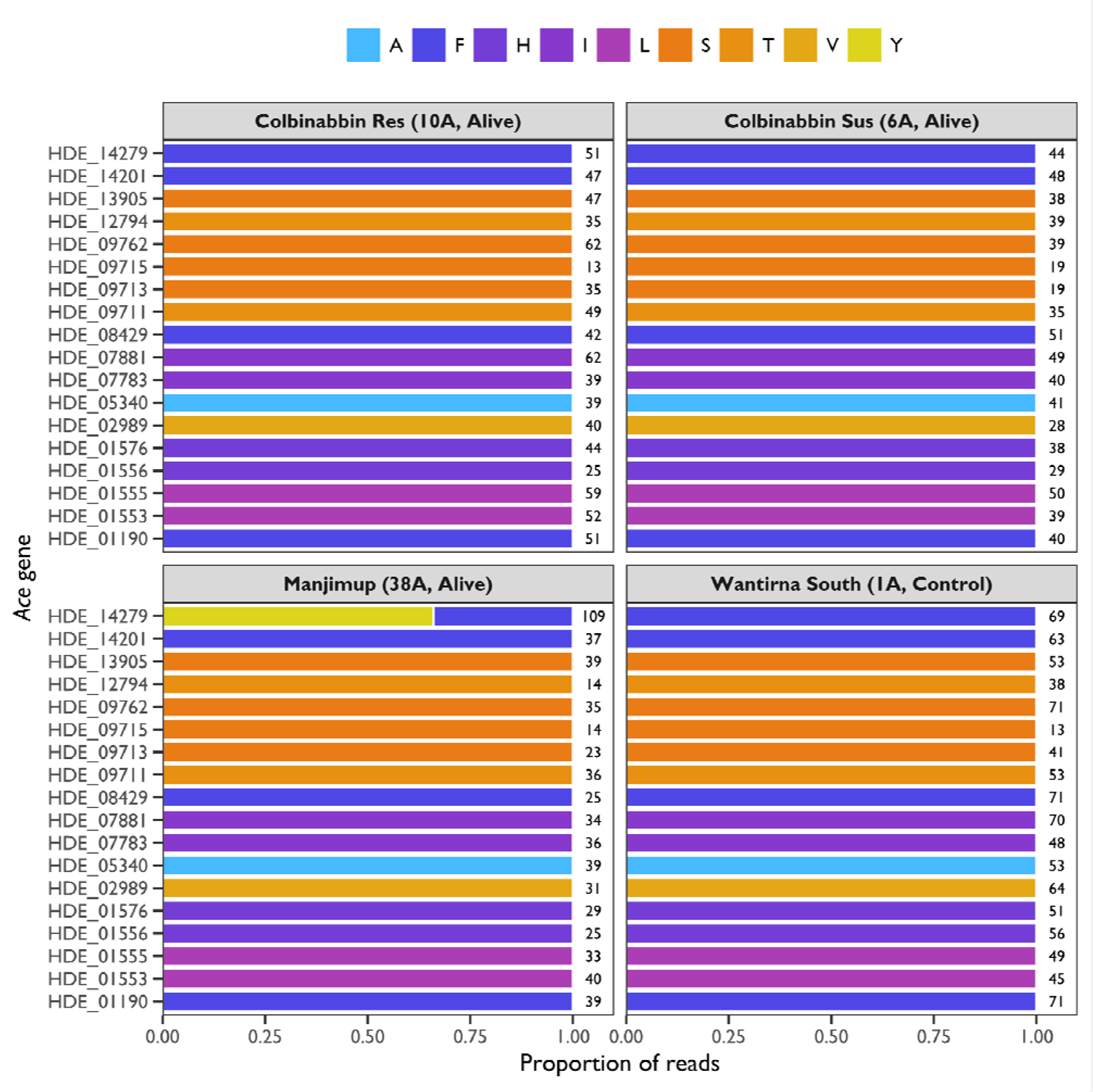
Proportion of reads for different amino acids at codon 331 (*Torpedo californica* numbering) across *ace* genes in *Halotydeus destructor*. The *x*-axis is the proportion of reads and the *y*-axis the *ace* gene. Horizontal bars represent the proportion of reads with nucleotide sequences encoding different amino acids, see legend. Numbers to the right of the bars indicate the number of observed reads that completely covered the codon. Paneels contain observations for select pool-seq libraries, read as ‘Population (DNA pool, Survival group)’. Note, Colbinabbin Res and Manjimup are resistant to organophosphates, whereas Colbinabbin Sus and Wantirna South are susceptible.

**Figure S6.**
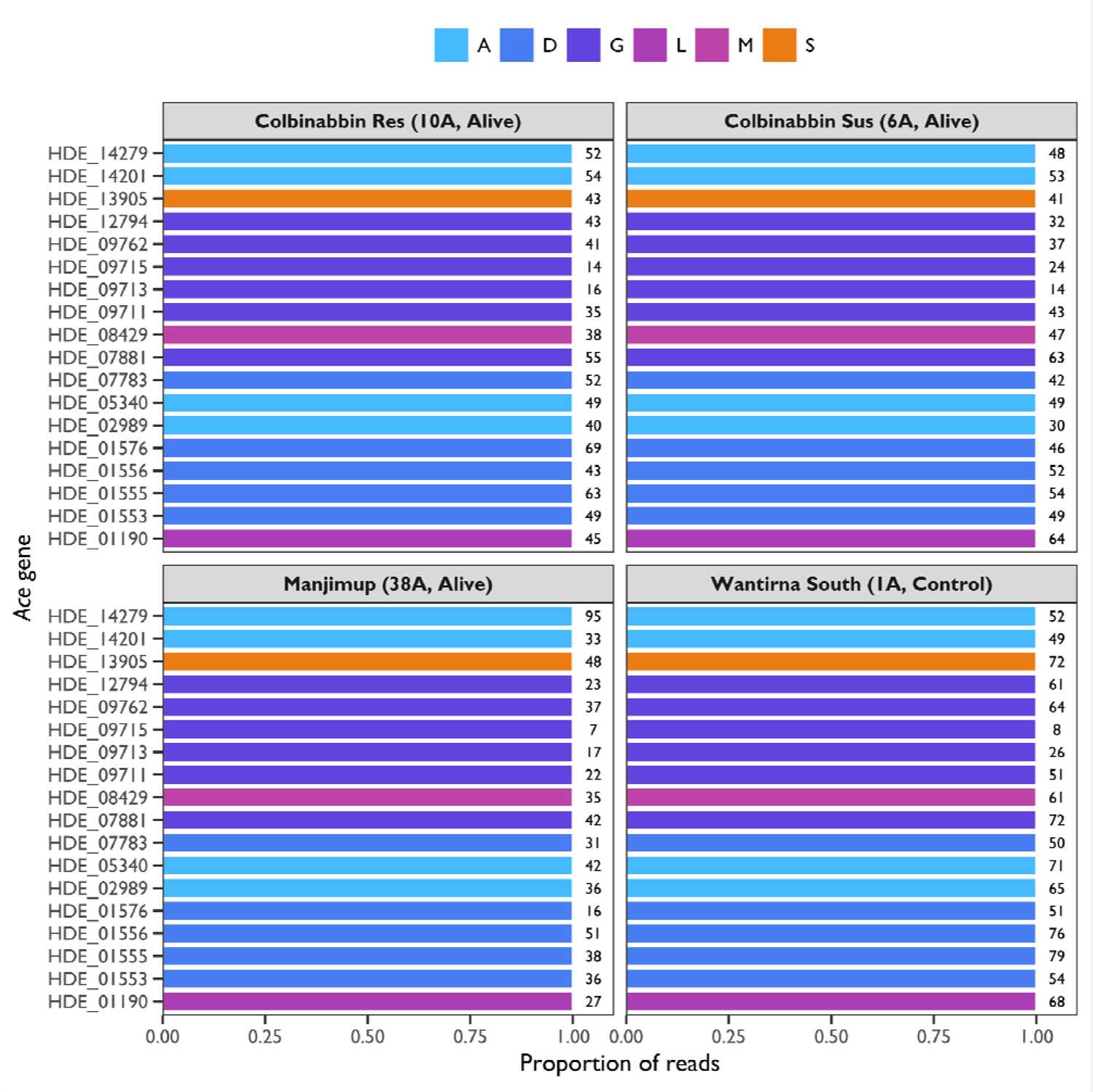
Proportion of reads for different amino acids at codon 441 (*Torpedo californica* numbering) across *ace* genes in *Halotydeus destructor*. The *x*-axis is the proportion of reads and the *y*-axis the *ace* gene. Horizontal bars represent the proportion of reads with nucleotide sequences encoding different amino acids, see legend. Numbers to the right of the bars indicate the number of observed reads that completely covered the codon. Paneels contain observations for select pool-seq libraries, read as ‘Population (DNA pool, Survival group)’. Note, Colbinabbin Res and Manjimup are resistant to organophosphates, whereas Colbinabbin Sus and Wantirna South are susceptible.

**Figure S7.**
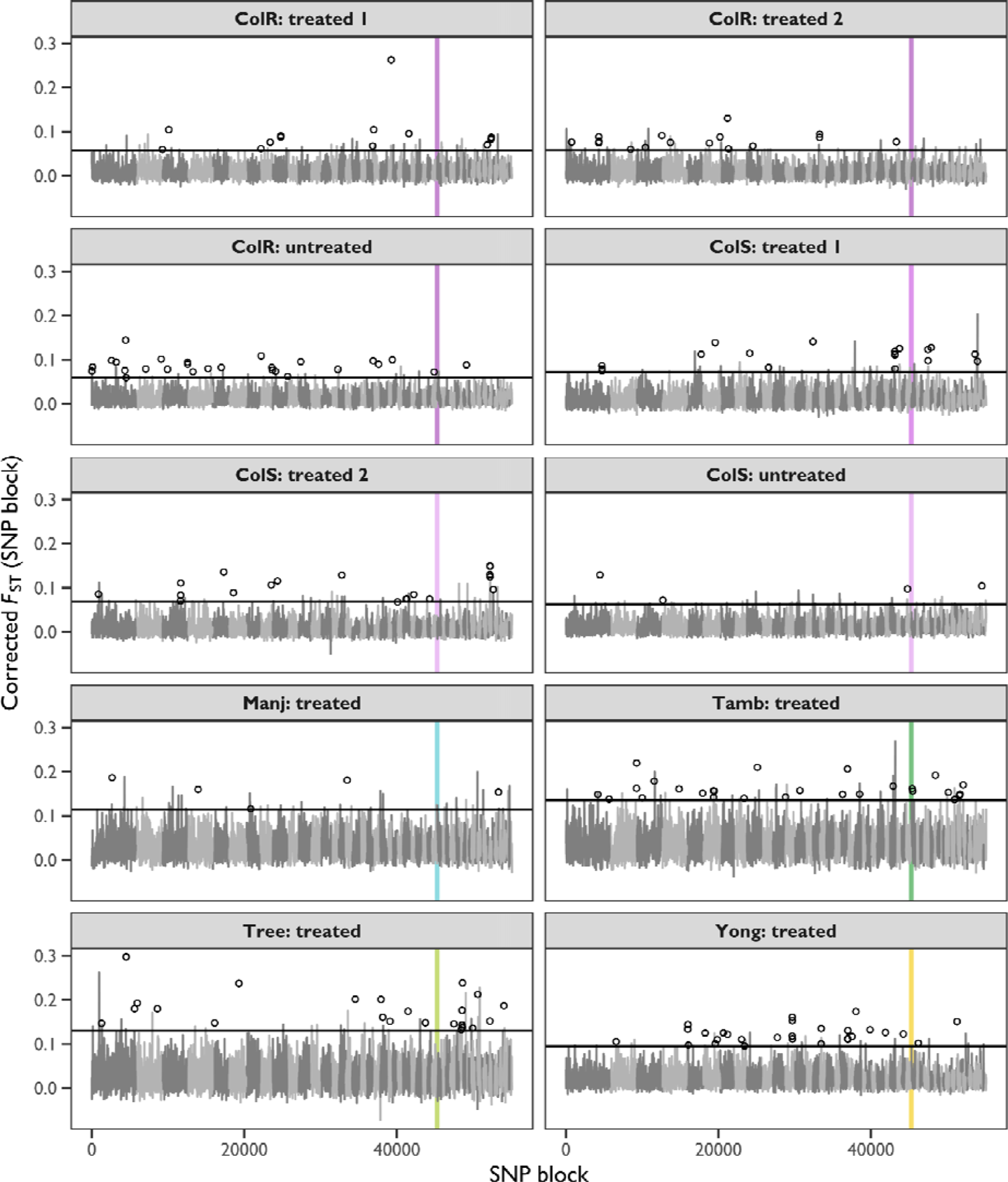
Genome outlier scans between alive and dead samples of *H. destructor* following organophosphate exposure. The *x*-axis represents the SNP block, indexed by contig size and linear order. The *y*-axis is the *F*_ST_ corrected for sampling varianc e. Black horizontal lines demarcate the 99^th^ percentile of *F*_ST_. Grey lines track the *F*_ST_ of *non-outlier* SNP blocks, with alternating shades of grey delimiting separate contigs. Open points denote *outlier* SNP blocks. Coloured vertical lines indicate SNP blocks containing the canonical *ace* gene. Panels contain different sample contrasts (see Table 1). Population codes are as follows: ‘ColR’ = Colbinabbin Res, ‘ColS’ = Colbinabbin Sus, ‘Manj’ = Manjimup, ‘Tamb’ = Tambellup, ‘Tree’ = Treeton, and ‘Yong’ = Yongarillup. Sample contrasts include ‘untreated’, which is the *F*_ST_ between two samples *unexposed* to organophosphate, and ‘treated’, which is the *F*_ST_ between samples of alive and dead mites that were *exposed* to organophosphate.

---

**Table S1.**
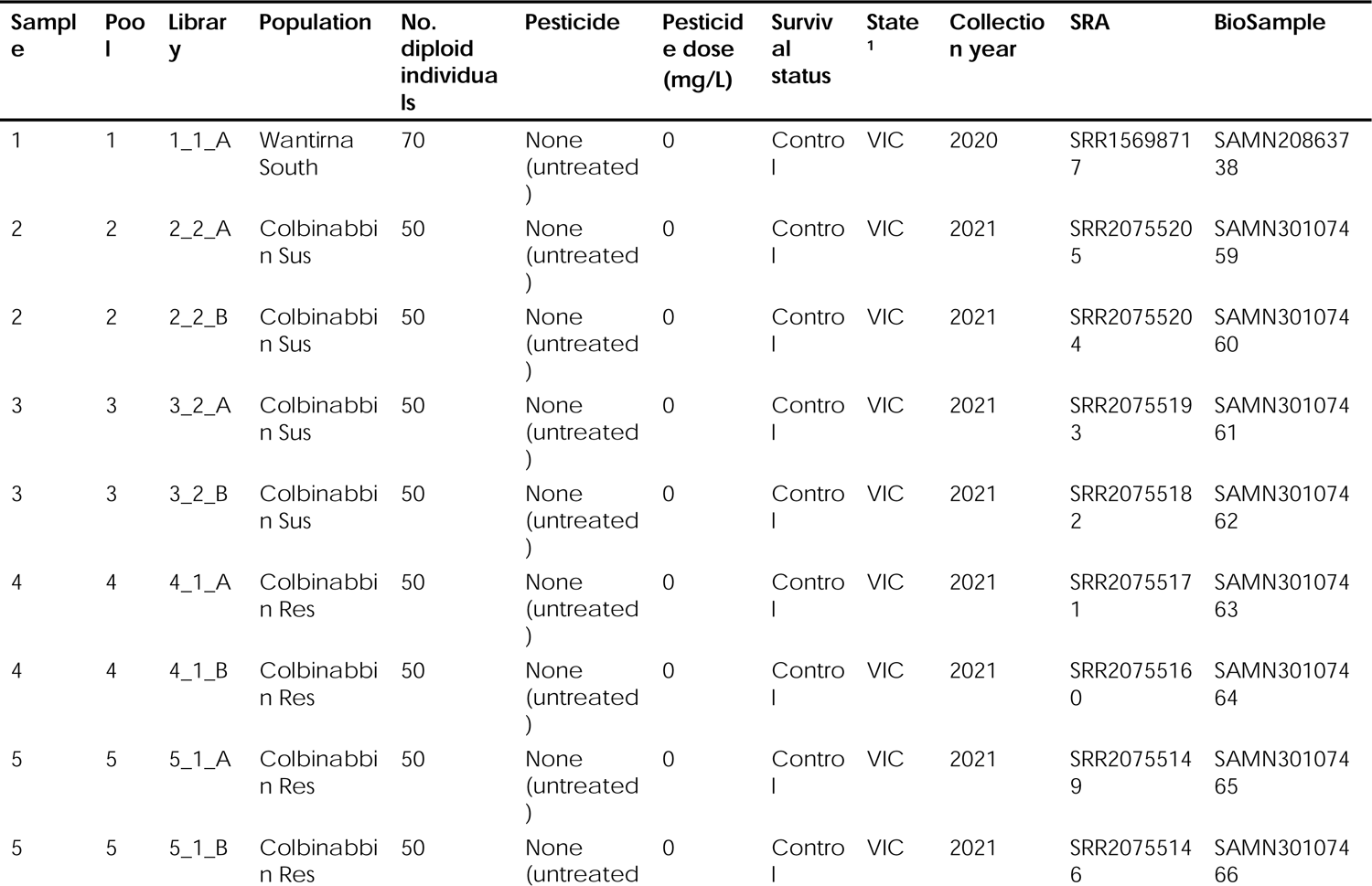
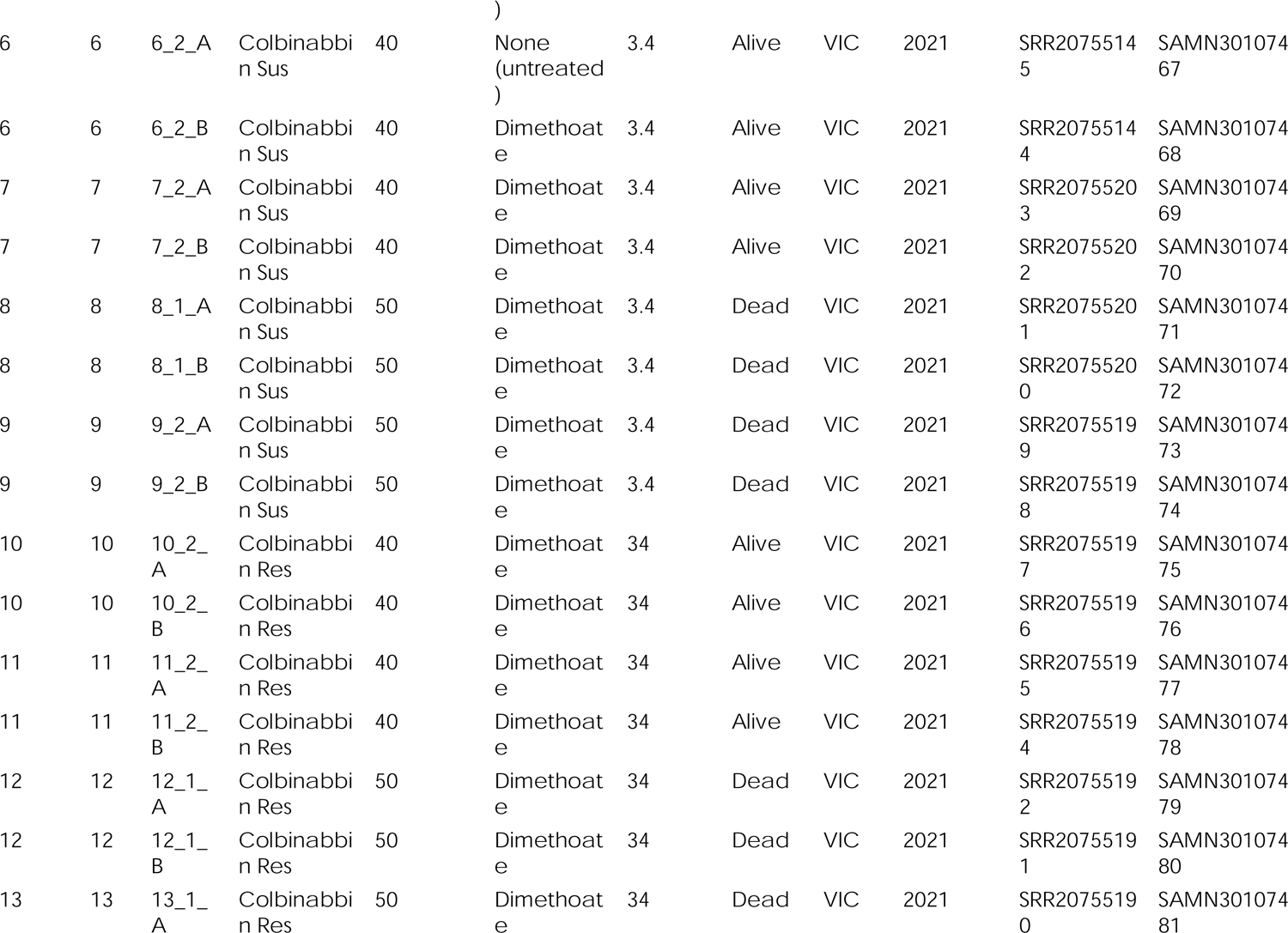
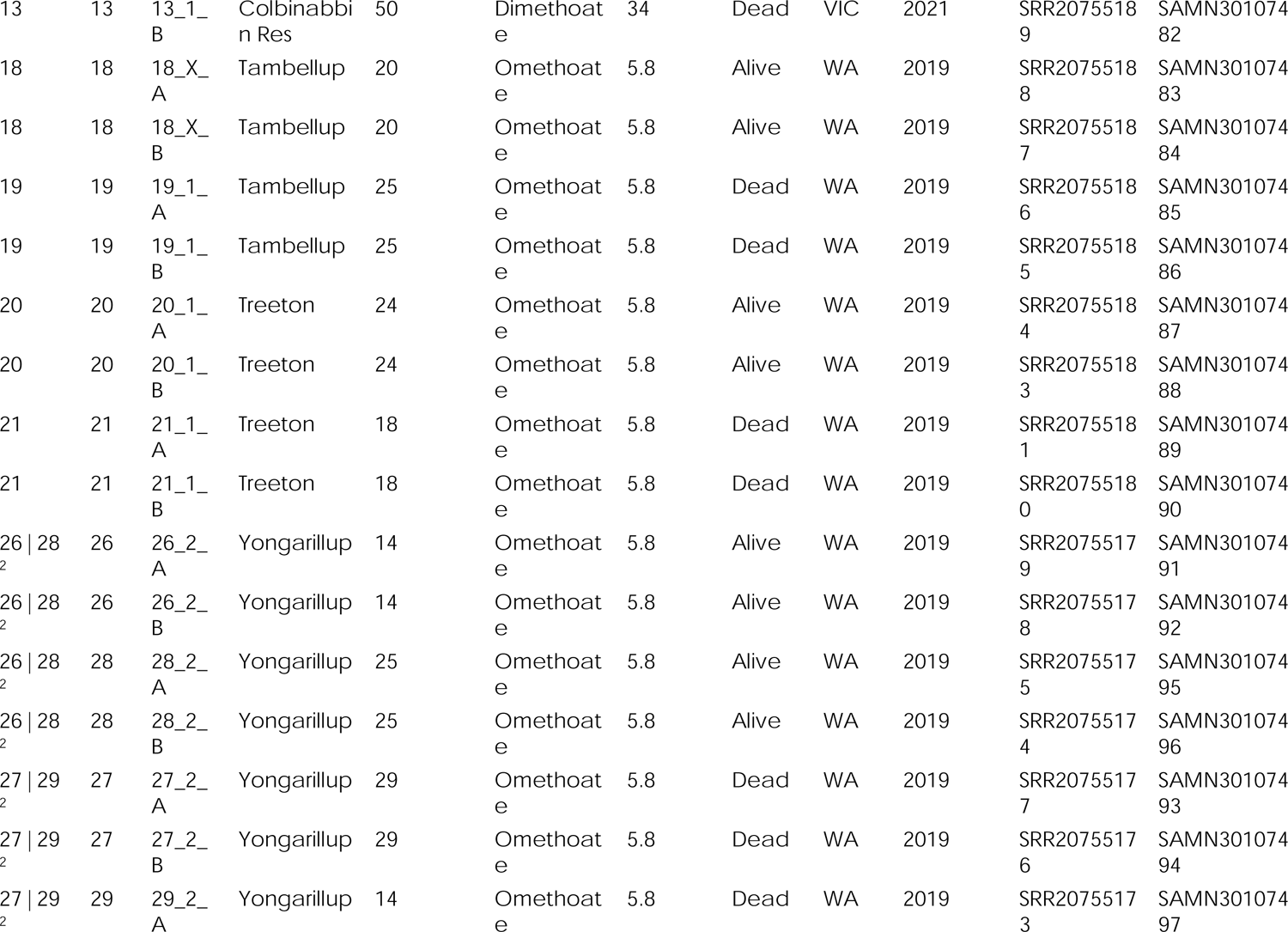
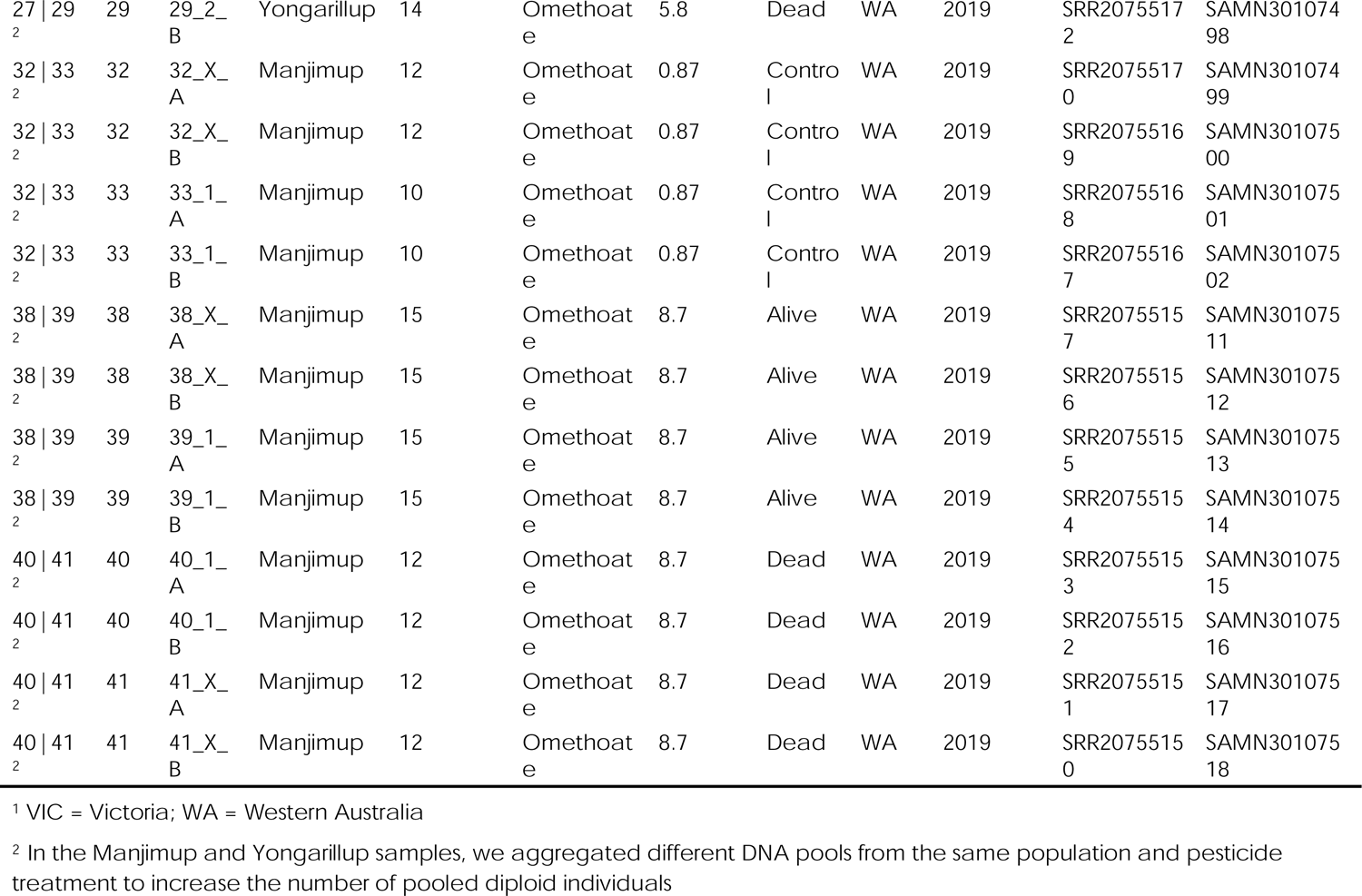
Metadata for pool-seq libraries. All *H. destructor* libraries were nested in DNA pools nested in biological samples (NCBI SRA and BioSample accession numbers listed for each library).

